# Genome-based discovery of pachysiphine synthases in *Tabernaemontana elegans*

**DOI:** 10.1101/2024.07.30.605783

**Authors:** Enzo Lezin, Mickael Durand, Caroline Birer Williams, Ana Luisa Lopez Vazquez, Thomas Perrot, Nicolas Gautron, Julien Pétrignet, Clément Cuello, Hans J. Jansen, Florent Magot, Sarah Szwarc, Pierre Le Pogam, Mehdi A. Beniddir, Konstantinos Koudounas, Audrey Oudin, Benoit St-Pierre, Nathalie Giglioli-Guivarc’h, Chao Sun, Nicolas Papon, Michael Krogh Jensen, Ron P. Dirks, Sarah E. O’Connor, Sébastien Besseau, Vincent Courdavault

**Affiliations:** Biomolécules et Biotechnologies Végétales, EA2106, Université de Tours, 37200 Tours, France; Laboratoire Synthèse et Isolement de Molécules BioActives (SIMBA, EA 7502), Université de Tours, 37200 Tours, France; Future Genomics Technologies, 2333 BE Leiden, The Netherlands; Équipe Chimie des Substances Naturelles, BioCIS, Université Paris-Saclay, CNRS, 91400 Orsay, France; Laboratory of Agricultural Chemistry, School of Agriculture, Aristotle University of Thessaloniki, 54124, Thessaloniki, Greece; Institute of Medicinal Plant Development, Chinese Academy of Medical Sciences and Peking Union Medical College, Beijing, China; Univ Angers, Univ Brest, IRF, SFR ICAT, F-49000 Angers, France; Novo Nordisk Foundation Center for Biosustainability, Technical University of Denmark, Kgs Lyngby, Denmark; Department of Natural Product Biosynthesis, Max Planck Institute for Chemical Ecology, Jena 07745, Germany

**Keywords:** pachysiphine, *Tabernaemontana elegans*, alkaloid, cytochrome P450, metabolic engineering, new-to-nature, P450 modeling

## Abstract

Plant specialized metabolism represents an inexhaustible source of active molecules, some of which have been used in human health for decades. Among these, monoterpene indole alkaloids (MIAs) include a wide range of valuable compounds with anticancer, antihypertensive, or neuroactive properties. This is particularly the case for the pachysiphine derivatives which show interesting antitumor and anti-alzheimer activities but accumulate at very low levels in several *Tabernaemontana* species. Unfortunately, genome data in *Tabernaemontanaceae* are lacking and knowledge on the biogenesis of pachysiphine-related MIAs *in planta* remains scarce, limiting the prospects for biotechnological supply of many pachysiphine-derived biopharmaceuticals. Here, we report a raw version of the toad tree (*Tabernaemontana elegans*) genome sequence. These new genomic resources led to the identification and characterization of a couple of genes encoding cytochrome P450 with pachysiphine synthase activity. Our phylogenomic and docking analyses highlights the different evolutionary processes that have been recruited to epoxidize the pachysiphine precursor tabersonine at a specific position and in a dedicated orientation, thus enriching our understanding of the diversification and speciation of the MIA metabolism in plants. These gene discoveries also allowed us to engineer the synthesis of MIAs in yeast through the combinatorial association of metabolic enzymes resulting in the tailor-made synthesis of non-natural MIAs. Overall, this work represents a step forward for the future supply of pachysiphine-derived drugs by microbial cell factories.

**Significance Statement:** While pachysiphine is a monoterpene indole alkaloid of high interest and the precursor of an anti-Alzheimer compound, its biosynthesis involving the epoxidation of tabersonine remains uncharacterized. By sequencing and assembling the genome of *Tabernaemontana elegans*, we identified two P450s exhibiting a pachysiphine synthase activity that we modelized to explore the evolutionary scenario leading to the acquisition of this expoxidase activity; and used to engineer yeast cell factories for securing pachysiphine supply and producing new-to-nature alkaloids.

## INTRODUCTION

Monoterpene indole alkaloids (MIAs) are a class of natural products mainly synthesized in plant species from the Gentianales order and in Cornales to a lower extent (St-Pierre *et al*., 2013). Besides encompassing complex chemistries, MIAs exhibit potent biological activities explaining their roles in the adaptation of plants to their environment and defense against bioagressors (Dugé De Bernonville *et al*., 2017). The marked toxicity of several MIAs has been notably valorized as pharmaceuticals to treat numerous human diseases as exemplified by the anticancer vinblastine/vincristine produced by the Madagascar periwinkle (*Catharanthus roseus*), the nootropic vincamine in *Vinca minor* or the antihypertensive ajmaline from *Rauvolfia* species. While knowledge on MIA biogenesis has continuously progressed since the 1970s, their supply remains limited by the restricted access to the medicinal source plants and by their low accumulation *in planta* (Courdavault *et al*., 2021). This impedes the exploitation of several MIAs, notably the promising anti-cancer and anti-alzheimer agent conophylline, which is synthesized in *Tabernaemontana* species (Umezawa and Shirabe, 2021; Zhou *et al*., 2023).

Due to their elaborate structures, MIAs stem from long and complex biosynthetic pathways endowed with distinct biosynthetic hubs that ensure the diversification of these compounds (Courdavault *et al*., 2014; Dugé de Bernonville *et al*., 2015; Kulagina *et al*., 2022). Strictosidine is the first and main hub from which almost all MIAs are derived. Indeed, the different decorations and/or cyclisations of the strictosidine terpenoid core lead to the formation of the three main MIA groups, including corynanthe, which encompasses tetrahydroalstonine and ajmalicine, iboga, with the famous catharanthine and ibogaine, and aspidosperma, such as the widely distributed tabersonine (O’Connor and Maresh, 2006).

Tabersonine stands out as a downstream MIA biosynthetic hub notably resulting in the synthesis of the vinblastine precursor vindoline in *C. roseus* and conophylline in *Tabernaemontana* species (Figure 1). The enzymatic modifications of tabersonine are indeed various, starting with P450-mediated hydroxylation at the C16 or C19 position, performed by tabersonine 16-hydroxylase (T16H; St-Pierre and De Luca, 1995; Besseau *et al*., 2013) and tabersonine 19-hydroxylase (T19H; Giddings *et al*., 2011), respectively. While 16-hydroxylation seems widespread in the aerial organs of *C. roseus*, 19-hydroxylation is restricted to the roots. Such a distribution results in the synthesis of different types of organ-specific MIAs. Indeed, the hydroxyl group at the C16 position can be *O*-methylated by 16-hydroxytabersonine 16-*O*-methyltransferase (16OMT) in the vindoline biosynthetic branch in leaves and flower (Levac *et al*., 2008; Lemos Cruz *et al*., 2023). By contrast, the hydroxyl group at the C19 position is acetylated by the region-specific tabersonine derivative 19-*O*-acetyltransferase to produce 19-acetyltabersonine (TAT; Carqueijeiro, Dugé De Bernonville, *et al*., 2018). Other hydroxylations of the tabersonine indole ring have been described, notably at the C15 and C17 positions, but the enzymes catalyzing these reactions have not yet been reported. Besides hydroxylation, tabersonine can also be oxygenated through epoxidation occurring at the C2-C3 and C6-C7 positions. In this respect, the cytochrome P450 tabersonine 3-oxygenase (T3O) converts the C2-C3 alkene of tabersonine into an epoxide in *C. roseus* leaves, which then spontaneously opens to form the corresponding imine alcohol (Kellner *et al*., 2015; Qu *et al*., 2015). This imine alcohol can be next reduced by the dehydrogenase/reductase tabersonine 3-reductase (T3R) to generate 2,3-dihydro-3-hydroxytabersonine that can be further *N*-methylated at the N1 position to initiate the synthesis of vindorosine (Besseau *et al*., 2013; Liscombe *et al*., 2010; Qu *et al*., 2015). Finally, the stereoselective C6-C7 epoxidation of tabersonine is ensured by tabersonine 6,7 epoxidase (TEX) to produce lochnericine in *C. roseus* roots, with the epoxide motif downward (α-epoxide; Carqueijeiro, Brown, *et al*., 2018). It should be noted that TEX cannot epoxidize 19-hydroxytabersonine whist 19-hydroxylation can follow TEX epoxidation for the synthesis of hörhammericine.

**Figure 1.**
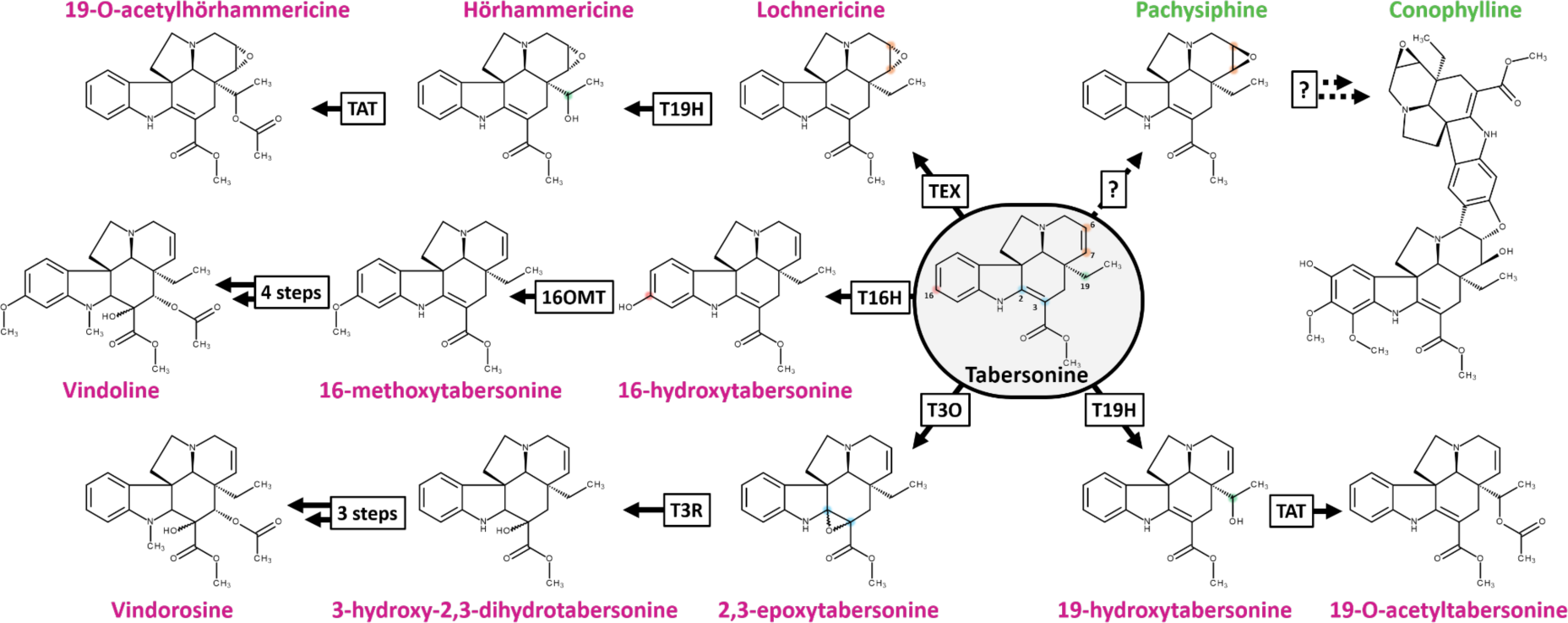
Tabersonine is a hub of the MIA metabolism. Characterized reactions from tabersonine are represented with solid arrows compared to dotted arrows for unknown reactions. Pink or green MIA names represent alkaloids from *Catharanthus roseus* or *Tabernaemontana elegans*, respectively. The position of each modified atom is annotated by a specific point color on the tabersonine and numbered with historical versions of numbering found in the enzymatic literature. *TEX: tabersonine epoxidase. T19H: tabersonine 19-hydroxylase. TAT: tabersonine acetyltransferase. T16H: tabersonine 16-hydroxylase. 16OMT: 16 O-methyltransferase. T3O: tabersonine 3-oxygenase. T3R: tabersonine 3-reductase*.

In addition to these decorations, tabersonine and its derivatives can be also self-condensed to form bis-indole alkaloids (Walia *et al*., 2020). Interestingly, the resulting dimers can engage tabersonine, various hydroxylated derivatives of tabersonine but also pachysiphine, a diastereoisomer of lochnericine with the epoxide upward (*β*-epoxide) (Walia *et al*., 2020). The MIA dimers formed with pachysiphine and its distinct hydroxy, methoxy and *N*-methyl derivatives notably include conophylline, conophyllidine, conophyllinine, conofoline, conolodinines A-D or melodinine K have been broadly reported in numerous *Tabernaemontana* species (Achenbach *et al*., 1994; Zèches-Hanrot *et al*., 1995; Kam *et al*., 2003; Kojima and Umezawa, 2006; Sim *et al*., 2019). Because it is strongly mobilized to fulfill the biosynthesis of conophylline and related compounds, pachysiphine is thus accumulated in very low amounts in *Tabernaemontana* species, preventing any prospects for the supply of potent conophylline derivatives through semi-synthetic approaches (Gonçalves *et al*., 2024). Most importantly, the enzymes involved in the metabolic pathway branch underlying the biosynthesis of conophylline derivatives from tabersonine in *Tabernaemontana* species remained unknown, a major obstacle to the development of metabolic engineering strategies for the production of these valuable natural products on an industrial scale (Figure 1).

The toad tree (*Tabernaemontana elegans*) is an important African medicinal plant that has attracted considerable attention since the early 1980s, and especially recently, for the elucidation of the MIA biosynthetic pathway (DeMars and O’Connor, 2024). The biotechnological potential *T. elegans* cell cultures have been particularly investigated as well as the pharmacological properties of its MIA extracts (Van Der Heijden *et al*., 1986; Van Der Heijden *et al*., 1989; Pallant *et al*., 2012; Mansoor *et al*., 2013; Ferreira and Paterna, 2019; Lucumi *et al*., 2002). The renewed interest in *T. elegans* for MIA metabolism has therefore prompted us to sequence the genome of this plant of interest and to exploit this valuable resource for the identification of MIA biosynthetic genes. In this work, we report a raw version of the *T. elegans* genome and the identification of two genes encoding P450s with pachysiphine synthase activity. These gene discoveries also allowed us to engineer the synthesis of MIAs in yeast through the combinatorial association of tabersonine-modifying enzymes resulting in the tailor-made synthesis of non-natural MIAs.

## RESULTS

### Genome sequencing and assembly of *Tabernaemontana elegans*

After extracting DNA from young leaves, we assembled the *T. elegans* genome using Flye (v2.8.3), followed by double polishing with Pilon (v1.2.3), leveraging both Oxford Nanopore Technologies (ONT) long-reads and Illumina short reads. The final assembly comprises 1,068,893,991 base pairs across 2,056 scaffolds, with an N50 of 4,500,278 base pairs for an L50 of 64 scaffolds, the longest of which spans 27,596,600 bp. The GC content of the assembled genome is 34.8%, which is consistent with expectations for genomes within Eudicotyledons. The genome assembly exhibits robust completeness, accuracy, and quality, with a k-mer completeness of 94.5% indicating comprehensive representation of raw reads, a quality value (QV) of 32.6 reflecting a base accuracy greater than 99.999%, and an LTR assembly index of 13.84, according to the reference genome criterion (Ou and Jiang, 2018), suggesting effective capture of repetitive elements. According to the identification of Eudicotyledon lineage using BUSCO, the assembled genome is 97.1% complete with a 2.8% low duplication rate (Table S1).

### Genome and transposon annotation

We constructed a *T. elegans* cDNA library by combining Illumina RNA sequencing of multiple plant organs (immature leaf, mature leaf, stem, flower bud, flower, and root) with a publicly available sequence read archive (SRR342013). By aligning the short reads to the assembled *T. elegans* genome, we annotated 25,045 protein encoding genes with high-confidence and a robust BUSCO completeness score of 97.0% (Table S1). In addition, we identified transposon sequences within the assembled genome, revealing significant proportions of different transposon types. The largest proportion of transposons belonged to the Long Terminal Repeat (LTR) class, which constituted 56.41% of the total genome. Terminal Inverted Repeat (TIR) transposons made up 7.93% of the genome, while non-Terminating Inverted Repeat (nonTIR) transposons accounted for 3.40%. (Table S2). Altogether, transposons comprised 67.73% of the assembled genome which is similar to the *Voacanga thouarsii* genome (Table S1, Figure 2B) (Cuello, Stander, Hans J Jansen, *et al*., 2022). This high percentage indicates a substantial presence of transposable elements within the genome, which could have important implications for its structural variation, evolution, and complexity, and may explain the low scaffolding levels reported here.

**Figure 2.**
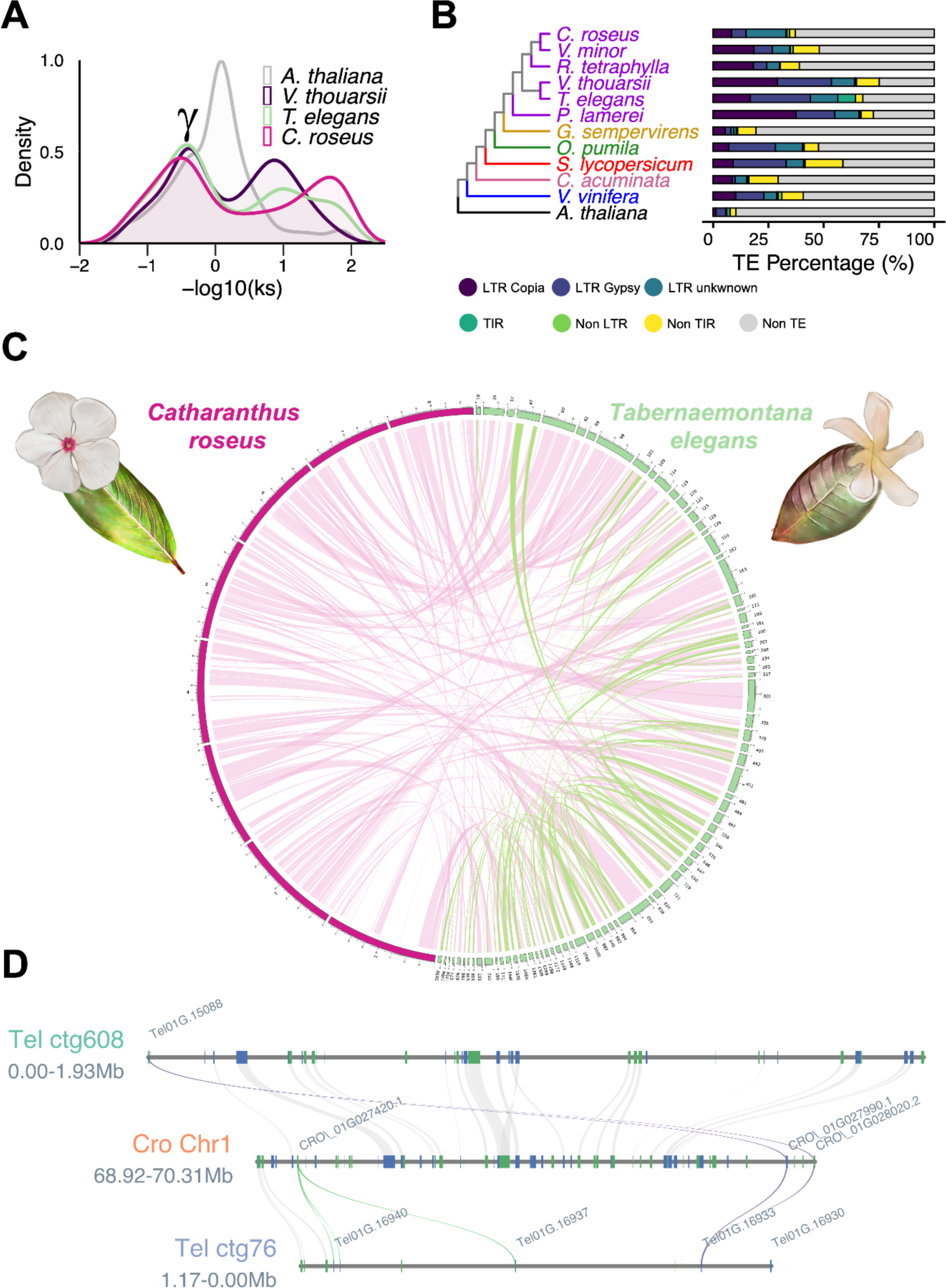
The annotated *T. elegans* genome. (**A**) Synonymous substitution rate (Ks) distribution plot for *T. elegans* paralogs compared to *A. thaliana*, *V. thouarsii* and *C. roseus*. γ indicates the conserved γ whole-genome triplication event that is shared among Eudicotyledons. (**B**) Phylogenetic tree of *T. elegans* and eleven other species, including five Apocynaceae (purple: *C. roseus*, *V. minor*, *R. tetraphylla*, *V. thouarsii* and *P. lamerei)*, one Gelsemiaceae (yellow: *G. sempervirens*), one Rubiaceae (green: *O. pumila*), one Solanaceae (red: *S. lycopersicum*), one Nyssaceae (pink: *C. acuminata*), one Vitaceae (blue: *V. vinifera*) and one Brassicaceae (black: *A. thaliana*). (**C**) Circos plot highlighting *T. elegans* genome-wide self-synteny (green links) and synteny between the *T. elegans* and *C. roseus* genomes (pink links). The eight *C. roseus* chromosomes are displayed in pink. Only the eighty *T. elegans* contigs containing eighty or more genes are shown in green. (**D**) Focus on the *T. elegans* genomic regions encompassing the PYS gene and the corresponding syntenic region retrieved in *C. roseus*. CYP71 (purple, PYS; green, other CYP71s) and gene synteny blocks are indicated by colored and gray lines, respectively.

### Whole genome duplication analysis

One of the major processes of plant diversification is whole genome duplication (WGD) (Landis *et al*., 2018). To investigate WGD in *T. elegans*, we determined paralogous gene pairs in twelve different species including three MIA-producing species from the Apocynaceae families (*C. roseus*, *V. thouarsii*, *T. elegans*), and one non-MIA-producing species (*A. thaliana*). We then determined the synonymous substitution per synonymous site (Ks) for each gene pair and presented it as -log10(Ks) (Figure 2A). For all species represented, the -log10(Ks) distribution clearly shows a major peak at approximately −0.4 (*i.e.* Ks ≈ 2.5), corresponding to the γ whole-genome triplication shared by Eudicotyledons (Jiao *et al*., 2012). Finally, the peaks identified at -log10(Ks) greater than 1 (i.e. Ks < 0,1) represent genes that are currently duplicated.

### Comparison of Orthologous genes

A maximum likelihood phylogenetic tree was generated by Orthofinder (Emms and Kelly, 2019) from 488 single-copy orthogroups, showing the apparent relationships between the twelve studied species (Figure 2B, Figure S1). Interestingly, the Apocynaceae species are divided into three groups. As shown by Cuello *et al*., (2024), *P. lamerei* represents an outgroup. The remaining species are divided into two distinct clades; one consisting of *C. roseus*, *V. minor* and *R. tetraphylla* (the Vinceae), and the other consisting of *V. thouarsii* and *T. elegans* (the Tabernaemontaneae). These results are consistent with existing molecular and morphological classifications (Yang *et al*., 2016; Simões *et al*., 2010).

### Synteny comparison

Intragenomic comparison of *T. elegans* reveals 147 synteny blocks comprising 1,213 gene pairs (Figure 2C, Table S3). Following this self-comparison, a genome-wide synteny analysis was performed between *T. elegans* and the closely related species *C. roseus*. Despite the fragmented nature of the *T. elegans* genome, the resulting circos plot clearly shows a high degree of conservation of the genomic organization, identifying 406 synteny blocks including 13,380 gene pairs (Figure 2C, Table S4). Furthermore, the synteny pattern is consistent with a 1:1 synteny ratio (Figure S2), as indicated by the Ks distribution (Figure 2A). Taken together, these data confirm the close evolutionary conservation between the *C. roseus* and *T. elegans* as reported in the previous section (Figure 2B).

### Predicting and identifying pachysiphine synthase candidates

Since pachysiphine derivatives and related dimers are widely distributed among *Tabernaemontana* species, we use our new *T. elegans* genome to initiate the search for the identification of a pachysiphine synthase. We thus undertook an homology-based search for candidates with TEX and T3O, the two already known enzymes epoxidizing tabersonine. We first focused on the nineteen closest TEX1 orthologous genes (≥50% identity), since pachysiphine has an epoxide group on the same C6-C7 position as lochnericine (Figure S3, Table S5). Fourteen of them were thus successfully amplified and cloned into the pHREAC expression vector before testing their activity (Tel01G.14148, Tel01G.22452, Tel01G.14149, Tel01G.21654, Tel01G.14146, Tel01G.11706, Tel01G.23700, Tel01G.23706, Tel01G.23689, Tel01G.21744, Tel01G.21725, Tel01G.21745, Tel01G.22828 and Tel01G.14838) (Figure 3A). Among the five remaining candidates, Tel01G.14841, Tel01G.9708, Tel01G.21647, Tel01G.21743 and Tel01G.23703 were too close to Tel01G.14838, Tel01G.21654, Tel01G.21744 and Tel01G.23706 to be distinguished by PCR and probably encode proteins with similar activity. The activity assay was performed by transient expression in *N. benthamiana* followed by incubation of the transformed leaf disks with tabersonine and LC-MS analysis of the reaction products (Lezin *et al*., 2023). A cursory analysis of each *m/z* ranging from 300 to 400 for the enzymatic product of each candidate, we observed no specific product formation for any of the candidates tested, except for Tel01G.21745 (Figure 3A). This candidate indeed synthesized a MIA with a retention time and MS/MS fragmentation similar to that of a pure 16-hydroxytabersonine standard thus strongly suggesting that it encodes a *bona fide* T16H (Figure S4). This result makes sense as this decoration potentially occurs in *T. elegans*. On the other hand, it strongly suggests that pachysiphine synthase does not belong to the TEX-like subfamily.

**Figure 3.**
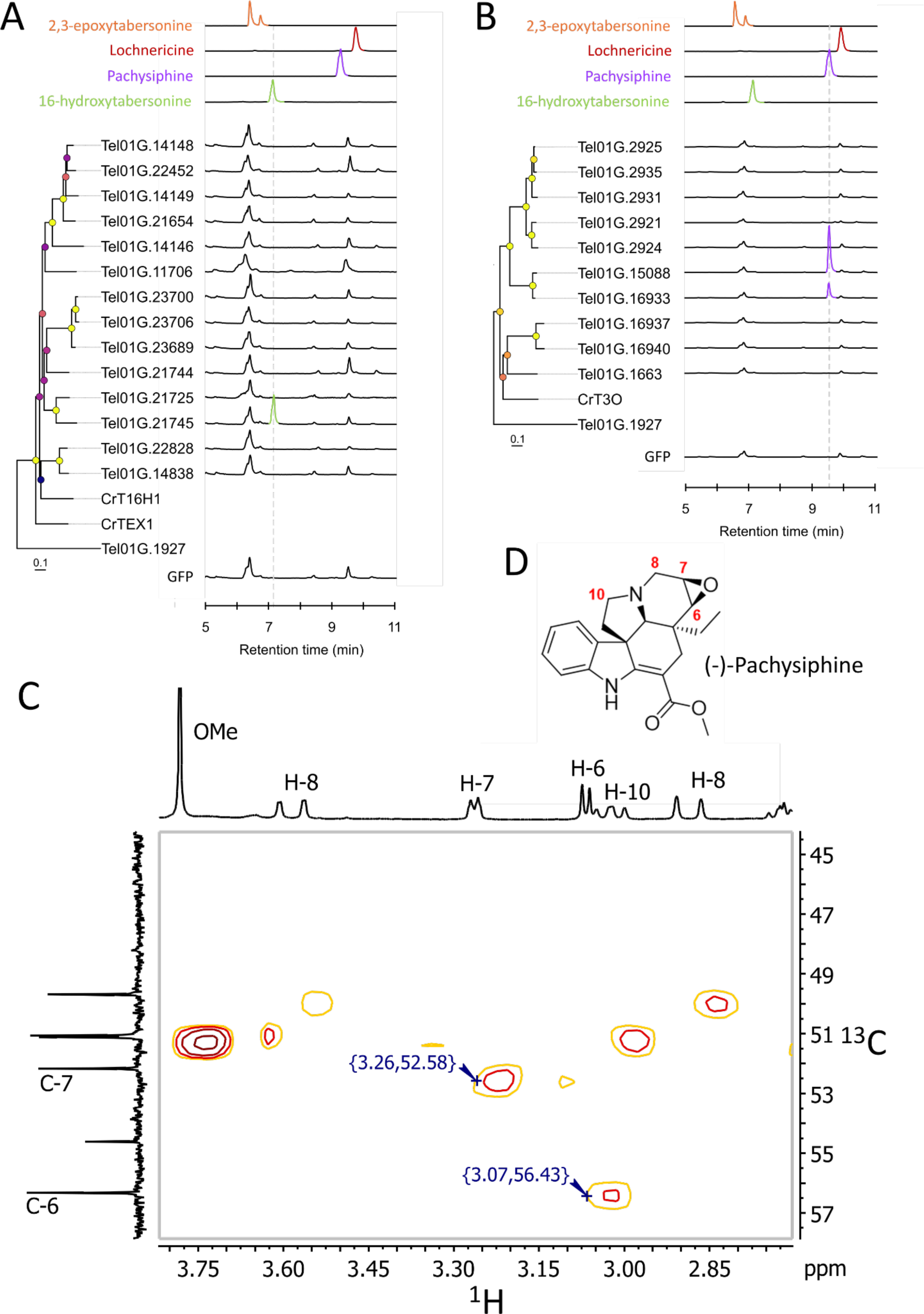
Screening of *T. elegans* P450s for oxygenation of tabersonine. TEX-like (**A**) and T3O-like (**B**) gene candidates and *GFP* were transiently overexpressed in tobacco leaves in the presence of tabersonine and reaction products were analyzed by LC-MS. Positive differential mass chromatograms of oxygenated tabersonine products at *m/z* 353 of P450 candidates are compared with GFP (negative control) and standards: 16-hydroxytabersonine (green line), pachysiphine (blue line), lochnericine (red line) and 2,3-epoxytabersonine (orange lin*e).* Metadata associated with bootstrap nodes are represented by a circle whose fill color scales proportionally with the bootstrap value (lower value, purple; higher value yellow), while branch lengths scale proportionally with substitutions per site value. Tel01G.1927 is the CYP71 P450 used to root the phylogenetic trees. Expanded HSQC spectrum (^1^H-^13^C correlation, focused on ^1^H 2.7-3.8 ppm and ^13^C 44-58 ppm) of the purified reaction product of Tel01G.15088 (**C**) characterized as (−)-pachysiphine (**D**). Full NMR data are available in Figure S7-S10. TEX, tabersonine-6,7-epoxydase; T3O, tabersonine 3-oxygenase; GFP, green fluorescent protein.

Based on this result, we next extended our candidate search to the more distant T3O-likes whose prominent member (T3O) epoxidizes tabersonine on the C2-C3 position. The ten closest T3O orthologous genes (Tel01G.2925, Tel01G.2935, Tel01G.2931, Tel01G.2921, Tel01G.2924, Tel01G.15088, Tel01G.16933, Tel01G.16937, Tel01G.16940, and Tel01G.1663) with high identity with T3O (≥50%) were selected and tested according to the previously described approach (Figure 3A, Figure S3, Table S5). Interestingly, while no product formation was observed at *m/z* 353 or any other *m/z* between 300 and 400 for 8 out of 10 candidates, a specific conversion of tabersonine into a compound with a retention time and a tandem mass spectrometry fragmentation profile similar to those of pachysiphine was observed for Tel01G.15088 and Tel01G.16933 (Figure 3B, Figure S5 and S6). Thus, although there is a slight difference in the amount of the MIA produced, this result suggests that both Tel01G.15088 and Tel01G.16933 encode pachysiphine synthases. To obtain firm evidence for the nature of their product, we carried out a milligram-scale production with Tel01G.15088 prior to purification and structure determination by NMR. The presence of the epoxide ring at the C6-C7 position was confirmed by the presence of signals resonating at 3.07 ppm and 3.26 ppm in the ^1^H NMR spectrum. Furthermore, the stereoselectivity of the epoxidation was confirmed by ^13^C NMR with respect to the C6 and C7 shifts (56.3 and 52.2 ppm), which are clearly different from those reported for lochnericine (57.4 and 54.1 ppm, respectively) (Figure 3C-D, Figure S7-S10) (Walia *et al*., 2020).

Taken together, these results thus definitively indicate that Tel01G.15088 and Tel01G.16933 encode two pachysiphine synthases (PYS) named PYS1 (GeneBank accession number XXXXXX) and PYS2 (GeneBank access number XXXXXX), respectively. Both PYS1 and PYS2 belong to the T3O-like subfamily of tabersonine metabolizing enzymes thus suggesting an evolutionary relationship between these enzymes and the T3O from *C. roseus*.

### Homology-based modeling differentiates tabersonine orientations in PYS and T3O

To gain further insights into the structural determinants leading to the evolution of PYS1/PYS2 and T3O activities, we undertook a homology-based modeling of these three enzymes resulting in the obtention of three models of very high confidence according to the calculated TM-scores of 0,918, 0,936 and 0,928 for PYS1, PYS2 and T3O, respectively (Figure S11A). Based on the crystal of CYP76AH1 from *Sativa miltiorrhiza* (Shi *et al*., 2021), the catalytic heme was placed in the active site and anchored to the conserved thiolate ligand (FXXGXXXCXXG) (Figure S11B) (Williams and De Luca, 2023). Using the three models, we next performed docking experiments with tabersonine in the proposed PYS1, PYS2 and T3O binding sites forming a pocket above the heme (Figure 4A-D). Tabersonine can thus be accommodated in each binding site with different poses. However, only one of them has the correct orientation and is close enough to the iron heme to allow tabersonine epoxidation at the correct site of metabolism (SOM) (i.e. C2-C3 for T3O and C6-C7 for PYS; Figure 4C-F). Furthermore, the best prediction for the tabersonine position inside PYS is not found in T3O and conversely. Consistent with the high sequence identity and similar activity of PYS1 and PYS2, the predicted models of both enzymes were highly similar, thus sharing identical binding pocket conformation and docking predictions, hence only PSY2 results are shown. However, comparing PYS2 and T3O revealed that 11 out of the 24 residues forming the substrate binding pocket were different (Figure 4G). Such a difference potentially affects the conformation of the PYS and T3O substrate binding sites and thus the orientation of tabersonine fixation. For instance, T3O has a large channel between the substrate binding site and the heme (Figure 4A-B). This directs the tabersonine towards the bottom of the substrate binding site thus favoring the interaction of the iron heme with the C2-C3 double bond of tabersonine (Figure 4C). In contrast, PYS2 has a reduced channel between the substrate binding site and the heme, which may limit the access to the iron heme to the only substrate moieties with a low steric hindrance such as the C6-C7 part of tabersonine (Figure 4D-E-F). These results therefore suggest that the two different channel conformations probably contribute to the positioning of tabersonine according to the more prone position for the required substrate site of metabolism (SOM). It is highly likely that this difference involves the substitution of two T3O residues by bulk residues in PYS1 and PYS2 (A374 to V373 and M378 to I377) as well as the insertion of a proline (P378) in these two enzymes (Figure 4B-E-G). Finally, docking predictions were used to identify the binding pocket residues potentially involved in the interactions with tabersonine (Figure 4H-I). In PYS2 and T3O, this fixation potentially involves the formation of hydrogen bonds between both the tabersonine methyl ester and amine functions and a serine residue (S122 in T3O and S211 in PYS2) (Figure 4G-I). Interestingly, it has to be noted that the S122/A123 and L210/S211 substitutions in the two enzymes redirect the position of the interacting serine residue and may also affect the orientation of tabersonine. In addition, the conserved T313/312 residue also participates in the formation of hydrogen bonds with the tabersonine methyl ester in PYS2 or the amine (Nx) in T3O. The specific orientations of tabersonine were also maintained by Pi-interactions with its aromatic cycle but also by van der Waals forces, both involving different residues in PYS and T3O (Figure 4H-I). For example, the I209/A210, A375/V373; M378/I377 and L490/I487 mutations in T3O and PYS2 may be crucial for the correct positioning of tabersonine in both types of binding pocket. All these results thus suggest that the different mutations in both the substrate binding sites and the channels probably explain how PYS2 and T3O evolve to acquire the capacity of epoxidizing tabersonine in two different positions.

**Figure 4.**
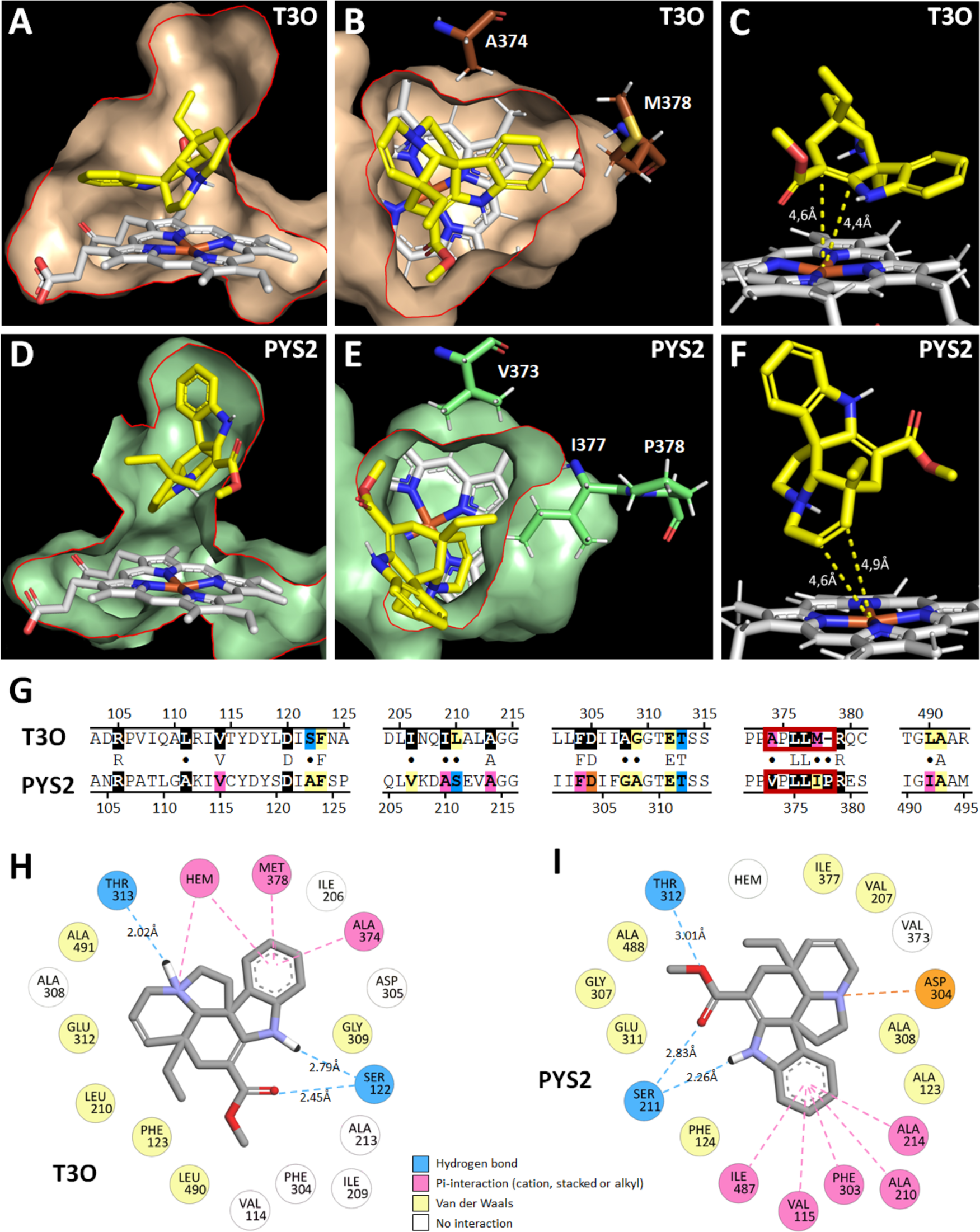
Molecular docking of tabersonine in PYS and T3O predicted structure. The results for PYS1 and PYS2 are similar, thus PYS2 (highest TM-score) is shown as the PYS enzymes model. The predicted substrate binding pocket of T3O (**A**, **B**) and PYS2 (**D**, **E**) containing the catalytic heme (in white) and the docked tabersonine (in yellow) are shown in longitudinal section (A,D) and cross-section (**B**, **E**) with a red line to underline the sectional plan. Tabersonine exposes the C2-C3 double bond to the iron heme in T3O (**C**) and C6-C7 double bond in PYS2 (**F**). T3O and PYS2 sequence alignment (**G**) of the five protein domains containing the 24 binding pocket residues (highlighted in color and black). Divergent residues between sequences are indicated by a dot. The red frame in the sequences indicates residues delimiting the channel between the substrate binding pocket and the heme (**B**, **E**). Residues interacting with tabersonine are indicated in a 2D diagram (**H**, **I**). Amino acids with equivalent position in the binding pocket of T30 (**H**) and PYS2 (**I**) are placed in the same order around tabersonine.

### Substrate specificity and subcellular localization of PYS1 and PYS2

Most of the P450s characterized to date are endoplasmic reticulum (ER)-anchored enzymes (Laursen *et al*., 2021). Consistent with this putative localization, both PYS1 and PYS2 bear a predicted membrane-spanning domain at the N-terminal end that may mediate ER anchoring (Figure S12). To confirm the role of these sequences, PYS1 and PYS2 were transiently expressed in *N. benthamiana* as yellow fluorescent protein (YFP) fusion proteins. The fusions were made exclusively at the YFP N-terminal end (PYS1-YFP and PYS2-YFP) to preserve the accessibility of the membrane-spanning domain and to avoid any anchoring interference. In epidermal cells of transformed *N. benthamiana* leaves, PYS1-YFP and PYS2-YFP exhibited a network-shaped fluorescent signal that perfectly merged with the signal of the ER cyan fluorescent protein (CFP) marker (Figure 5A-H). Thus, this result strongly suggests that both PYS1 and PYS2 are anchored to the ER, in agreement with the previously reported localization of P450s related to MIA synthesis and notably T3O (Guirimand *et al*., 2009; Besseau *et al*., 2013; Dugé de Bernonville *et al*., 2015; Kellner *et al*., 2015; Parage *et al*., 2016; Tatsis *et al*., 2017).

**Figure 5.**
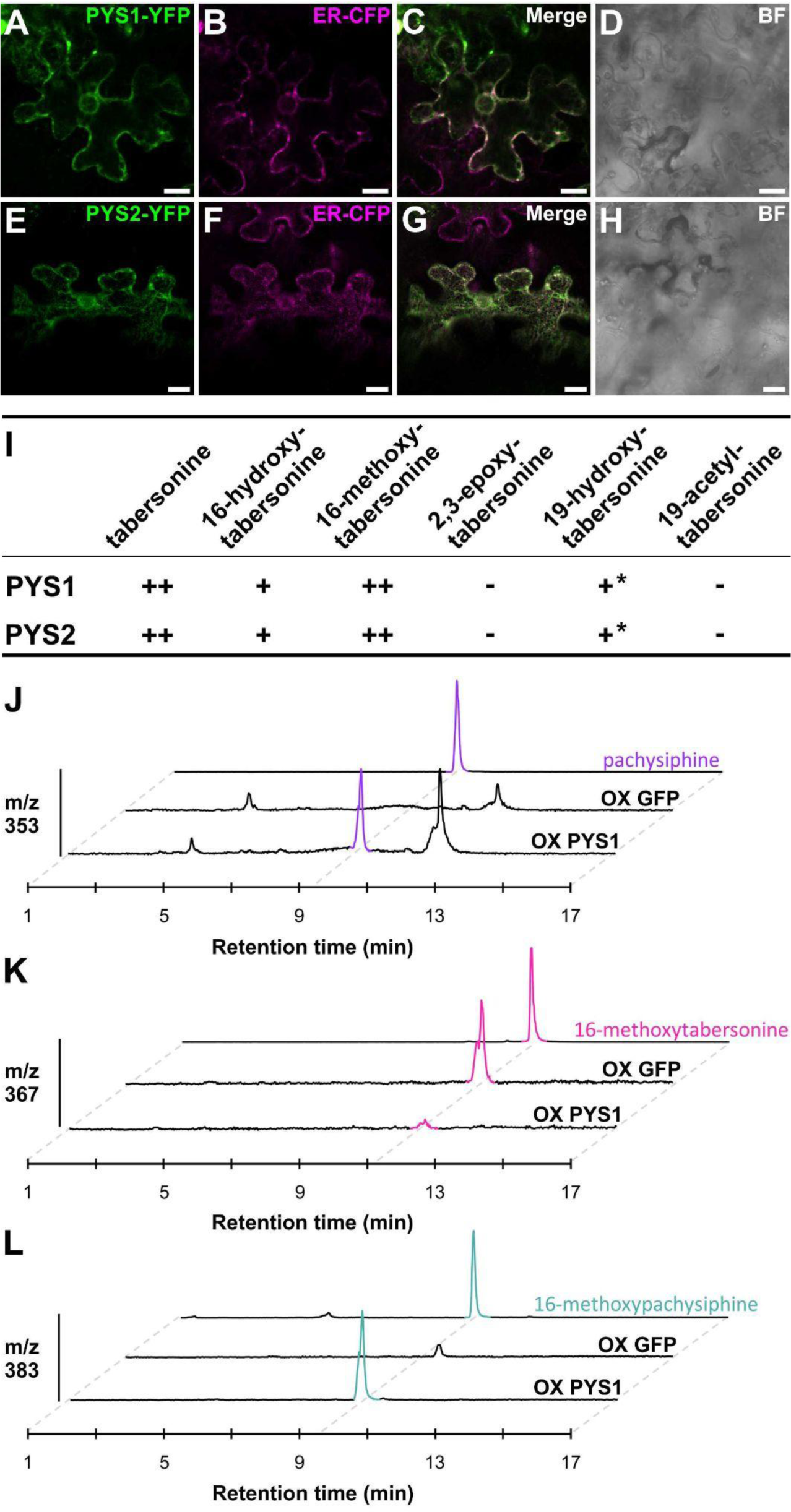
Subcellular localization and substrate uptake of PYS1 and PYS2. Epidermal cells of N. benthamiana were transiently transformed with the PYS1-YFP (**A**) or PYS2-YFP construct (**B**) together with an ER-CFP marker (**B**, **F**). Co-localization was confirmed after merging the two fluorescence signals (**C**, **G**). Cell morphology (**D**, **H**) was observed by bright field (BF). Bars = 10μm. (**I**) Activity of PYS1 and PYS2 with different tabersonine derivatives. PYS1 and PYS2 were transiently overexpressed in tobacco leaves in the presence of tabersonine derivatives. The reaction products were analyzed by LC-MS to determine the PYS activity. (**J**-**L**) *PYS1* or *GFP* genes were transiently overexpressed in *Catharanthus roseus* young leaves to analyze the *Catharanthus* MIA metabolism by LC-MS. Both conditions were studied in three plant replicates but one representative replicate for each one was shown. Positive differential mass chromatograms were analyzed at *m/z* 353 (**J**), *m/z* 367 (**K**) and *m/z* 383 (**L**) corresponding to [M+H]^+^ of pachysiphine (purple color), 16-methoxytabersonine (pink color) and 16-methoxypachysiphine (blue color) standards, respectively. MS/MS of pachysiphine and 16-methoxypachysiphine produced in *C. roseus* leaves were compared to standards in Figure S14A-B. At *m/z* 337, tabersonine is barely detectable in negative controls (Figure S14C). *++: high activity. +: low activity. -: no activity. *: low activity in tobacco leaves but no activity in yeast. PYS: pachysiphine synthase. OX: overexpression. m/z: mass to charge ratio. LC-MS: liquid chromatography coupled to mass spectrometry*.

To gain more insight into the biochemical properties of PYS1 and PYS2, both enzymes were assayed with all available tabersonine derivatives including 16-hydroxy and 16-methoxytabersonine, 19-hydroxy and 19-acetyltabersonine, and 2,3-epoxytabersonine. PYS1 and PYS2 were thus transiently or stably expressed in *N. benthamiana* and *S. cerevisiae*, respectively, before being fed with the different substrates for 48h (Figure 5I, Figure S13). In both heterologous organisms, similar substrate acceptances were monitored for tabersonine derivatives decorated on the indole ring (16-hydroxy and 16-methoxytabersonine) while the C2-C3 epoxidation completely inhibited PYS1 and PYS2 activity. On the other hand, notable/slight differences in the epoxidation activity were observed when the modification occurred on the tabersonine ethyl group. Indeed, while 19-hydroxylation and 19-acetylation of tabersonine prevent the C6-C7 epoxidation catalyzed by PYS1 and PYS2 in yeast, we found that both enzymes were capable of epoxidizing 19-hydroxytabersonine at low levels when expressed in *N. benthamiana*. Overall, these substrate acceptances are similar to those of TEX and may be related to the diversity of MIAs derived from pachysiphine. Interestingly, although no 19-hydroxy or 19-acetyl derivatives of pachysiphine have been described so far, the plasticity of the substrate acceptances of PYS1 and PYS2 towards these substrates opens up new possibilities for the synthesis of new-to-nature MIAs.

Finally, to obtain additional evidence of PYS activity, we performed a functional assay *in planta*. Since no functional genomic tool has been developed for *T. elegans* and, and in particular no gene silencing procedure, we transiently overexpressed PYS1 in the leaves of *C. roseus*, which do not synthesize pachysiphine or derived MIAs. Interestingly, compared to leaves overexpressing GFP, PYS1 overexpression resulted in the accumulation of pachysiphine, according to the similarity of retention time and MS/MS spectra with a pure standard (Figure 5J, Figure S14A-B). This was also accompanied by the consumption of tabersonine, which accumulated at a low amount in the GFP-overexpression plants (Figure S14C). In addition, we also observed the accumulation of 16methoxypachysiphine together with a correlated consumption of 16-hydroxypachysiphine as a direct consequence of the redirection of metabolic fluxes (Figure 5K-L). All together, these results thus strongly suggest that PYS1 exhibits an efficient pachysiphine synthase activity *in planta*.

### Insight into the genomic context and evolution of PYS1 and PYS2

Since PYS1 and PYS2 share a common ancestor with T3O (Figure S3), we sought to gain deeper insight into the evolutionary origins of PYS. Interestingly, the two *T. elegans* genomic regions encompassing *PYS1* and *PYS2* exhibit synteny with chromosome 1 of *C. roseus* (Crov3, Li *et al*., 2023) (Figure 2D). Remarkably, the *C. roseus* syntenic region is flanked by *CYP71* genes including *CYP71D1* (=CRO_01G028020.2) and *CYP71D2* (=CRO_01G027990.1) (Miettinen *et al*., 2014). On the one hand, *CYP71D1* and *CYP71D2* share high sequence homology with *PYS*, while on the other hand, *CRO_01G027420.2* shares high sequence homology with *Tel01G.16937* and *Tel.01G16940*. The neighbor-joining phylogenetic tree clearly shows that Tel01G.16937 and Tel.01G16940 are orthologs of CRO_01G027420.2 which is a paralog of T3O (Figure S15). Despite this, these two protein-coding genes do not exhibit oxygenase activity on tabersonine (Figure 3B). In addition, PYS1 and PYS2 are clearly orthologous to CYP71D1 and CYP71D2 in *T. elegans,* suggesting an evolutionary relationship. Interestingly, the *T. elegans* syntenic fragment containing PYS2 not only hosts PYS2 paralogs (*i.e.* Tel01G.16937 and Tel01G.16940) but also includes a putative *N*-methyltransferase (NMT) encoding gene (*Tel01G.16930*). This is a relevant feature since NMTs are MIA tailoring enzymes and this gene may therefore be involved in MIA biosynthesis.

### Identification of *N*- and *O*-methyltransferases of pachysiphine and related MIA

To gain insight into the potential role of the *T. elegans* NMT encoding gene in MIA metabolism, its coding sequence was amplified and cloned into the pHREAC vector for functional assays. However, no *N*-methylation of tabersonine and pachysiphine were observed in our experimental conditions thus suggesting that Tel01G.16930 is not active on these substrates (Figure 6A). The close vicinity of this putative NMT and PYS2 in the *T. elegans* genome also prompted us to investigate the genomic environment of TeT16H. In *C. roseus*, the genes encoding the two contiguous T16H1 and T16H2 are indeed adjacent to *16OMT1* (Cuello, Stander, Hans J. Jansen, *et al*., 2022; Li *et al*., 2023). However, no gene encoding enzymes displaying methyltransferase features have been identified in the vicinity of TeT16H. We therefore performed a homology-based search using *C. roseus* 16OMT1 as a bait to identify candidates for 16-hydroxytabersonine and 16-hydroxypachysiphine *O*-methyltransferases in the entire *T. elegans* genome. A set of 10 genes with high identity to 16OMT1 (Tel01G.8583, Tel01G.8582, Tel01G.8584, Tel01G.8589, Tel01G.6570, Tel01G.21576, Tel01G.21570, Tel01G.21569) was retrieved and each candidate was functionally tested by transient expression in *N. benthamiana* together with Tel01G.21745 (TeT16H) and feeding with tabersonine. While no *O*-methylation occurred in the leaves expressing GFP, Tel01G.8589, Tel01G.6570, Tel01G.21576, Tel01G.21570 and Tel01G.21569, the formation of a compound with a retention time and a mass fragmentation profile similar to those of 16-methoxytabersonine was observed with Tel01G.8582, Tel01G.8583 and Tel01G.8584 (Figure 6B, Figures S16, S17 and S18). Interestingly, we noted that these three enzymes were also able to *O*-methylate 16-hydroxypachysiphine (Figure S19). All these results thus suggest that Tel01G.8582, Tel01G.8583 and Tel01G.8584 encode 16-hydroxytabersonine/16-hydroxypachysiphine 16-*O*-methyltransferases, which we have named Te16OMT1, Te16OMT2 and Te16OMT3, respectively. Furthermore, these three genes are close neighbors in the *T. elegans* genome suggesting that both genes arose from a local duplication event.

**Figure 6:**
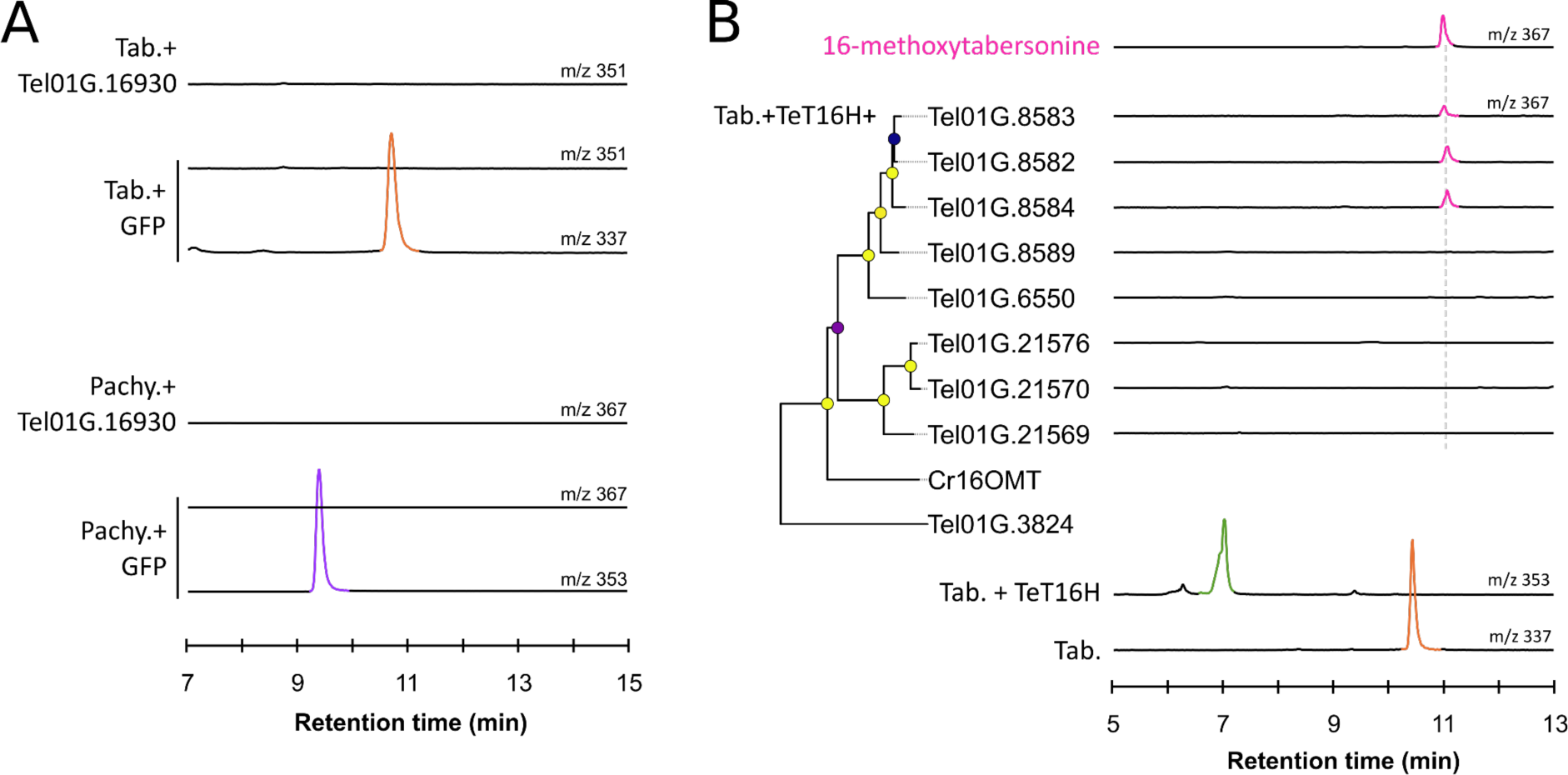
Screening of methyltransferase activities for methylation of tabersonine derivatives. (**A**) Screening of *N*-methyltransferase activity with tabersonine and pachysiphine. Positive differential mass chromatograms of tabersonine at m/z 337, putative N-methyltabersonine at m/z 351, pachysiphine at m/z 353 and putative N-methylpachysiphine at m/z 367. The NMT gene candidate, Tel01G.16930, was transiently overexpressed in tobacco leaves in the presence of tabersonine or pachysiphine and reaction products were analyzed by LC-MS and compared to a GFP negative control. (**B**) Screening of O-Methyl-Transferase activities with 16-hydroxytabersonine produced by the TeT16H from tabersonine. OMT candidates are arranged according to neighbor-joining phylogenetic tree classification (100 bootstrap replicates). Metadata associated with bootstrap nodes are represented by a circle whose fill color scale proportionally with the bootstrap value (lower value, purple; higher value yellow) while branch lengths scales proportionally with substitutions per site value. The scale bar represents 0.1 substitutions per site. Positive differential mass chromatograms of tabersonine at *m/z* 337, 16-hydroxylated tabersonine at *m/z* 353 and the *O*-methyltransferase assays with OMTs 16OMT-like candidates at *m/z* 367 compared to a 16-methoxytabersonine standard (pink line). OMT gene candidates and *TeT16H* were transiently co-overexpressed in tobacco leaves in the presence of tabersonine and reaction products were analyzed by LC-MS and compared to a GFP negative control. *m/z: mass to charge ratio. Tab.: tabersonine. pachy.: pachysiphine. TeT16H: tabersonine-16-hydroxylase from T. elegans. OMT: O-methyltransferase. GFP: green fluorescent protein. LC-MS: liquid chromatography coupled to mass spectrometry*.

### Metabolic engineering of the pachysiphine biosynthesis

Due to promising pharmacological properties of pachysiphine and related compounds, we developed a metabolic approach based on tabersonine bioconversion to explore the controlled synthesis of new-to-nature tabersonine derivatives. Therefore, we co-expressed in yeast a combination of tabersonine metabolizing enzymes including T16H, 16OMT, T19H and TAT from *C. roseus* besides PYS1. Based on the PYS1 substrate acceptance and its incapacity to utilize 19-hydroxytabersonine, we engineered two distinct yeast cell factories. Both yeast strains were successively cultured and the establishment of a synthetic biosynthetic route was followed by the detection of the different biosynthetic intermediates generated along the bioconversion assays. The first strain expressed the *C. roseus* T16H1 and 16OMT1, as well as PYS1 (TOP strain) while the second strain integrated both the *C. roseus* T19H and TAT (TT strain) (Figure 7). The TOP strain was cultured for 48 hours in a rich medium (YPD) containing 40 µM of tabersonine, resulting in the production of a compound with a [M+H]^+^ of 383 as expected for 16-methoxypachysiphine. Interestingly, this metabolite, as well as all other MIAs produced in yeast, were secreted into the yeast culture media (Kulagina *et al*., 2022). The TOP yeast cells were then removed by centrifugation and the collected supernatant was then inoculated with the TT strain. After a further 24 hours of culture, the potential 16-methoxypachysiphine completely disappeared and we saw the formation of another compound at [M+H]^+^ of 441 in the culture medium suggesting the synthesis of 16-methoxy-19-acetylpachysiphine. This result clearly demonstrates that the combination of enzymes from different plants allows the tailor-made synthesis of new-to-nature tabersonine/pachysiphine derivatives in yeast heterologous systems.

**Figure 7.**
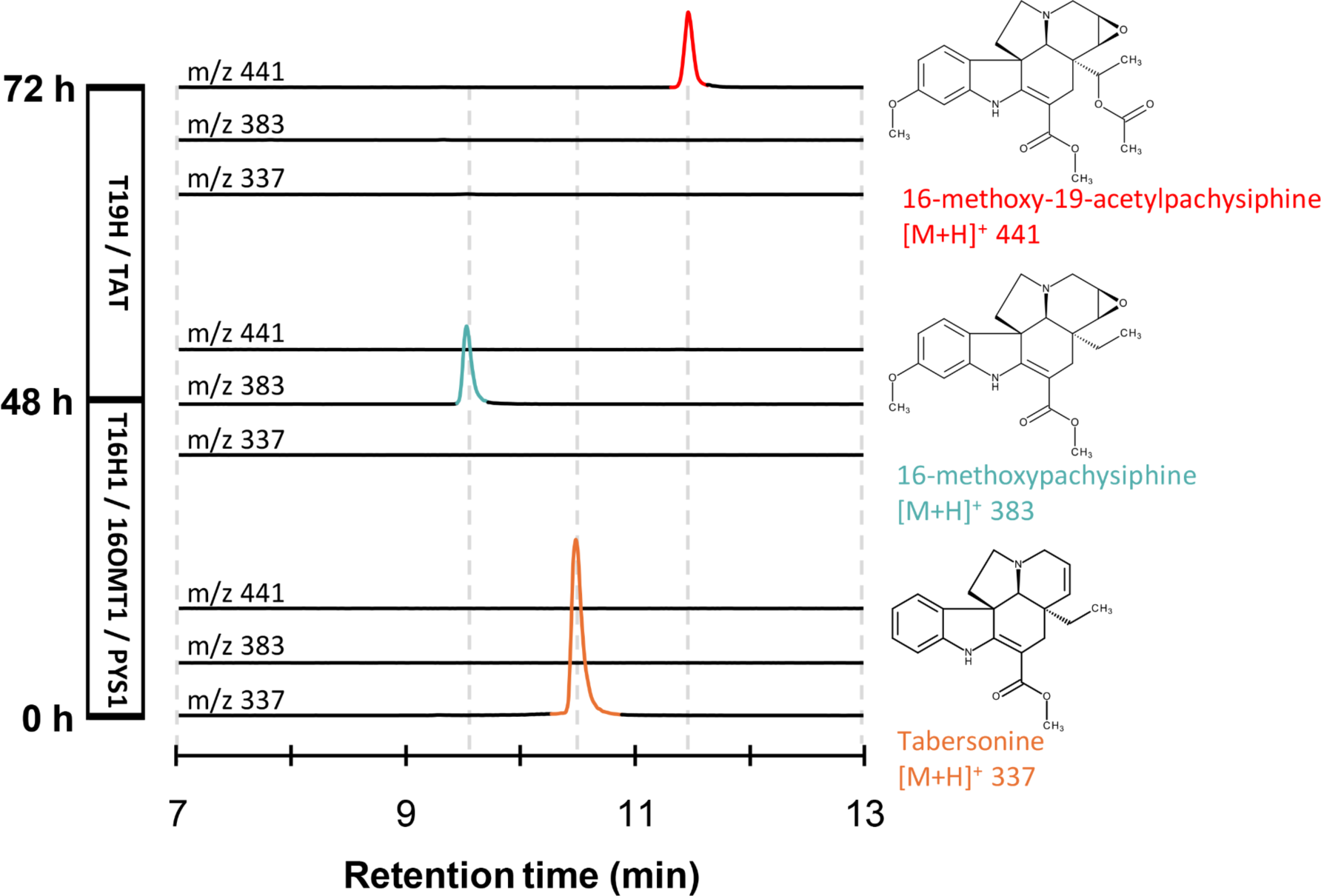
Biosynthesis of a new-to-nature MIA with PYS1 in yeast. Yeast expressing T16H1, 16OMT1 and PYS1, named “TOP”, was fed with tabersonine (lower chromatograms, at *m/z* 337, in orange) and grown for 48 h to produce 16-methoxypachysiphine (middle chromatograms, at *m/z* 383, in blue). The “TOP” strain was removed and a yeast expressing T19H and TAT, named “TT”, was added to “TOP” resulting enzymatic products. The “TT” strain was grown for 24 h, resulting in the production of 16-methoxy-19-acetylpachysiphine production (top chromatograms, at m/z 441, in red). The three consecutive positive differential mass chromatograms at *m/z* 337, *m/z* 383 and *m/z* 441 show, respectively, the substrate and the resulting MIAs produced by the “TOP” and “TT’’ strains in the yeast culture medium at 0, 48 and 72 h after tabersonine feeding. *m/z: mass-to-charge ratio. T16H1: tabersonine-16-hydroxylase 1. 16OMT1: 16-O-methyltransferase 1. PYS1: pachysiphine synthase 1. T19H: tabersonine 19-hydroxylase. TAT: tabersonine acetyltransferase. LC-MS: liquid chromatography coupled to mass spectrometry*.

## DISCUSSION

Although *T. elegans* is an Apocynaceae species of prime importance regarding the elucidation of the MIA chemical diversity and metabolism, no genomic data were yet available for this plant. This prompted us to sequence the genome of this medicinal plant in order to enrich the dataset dedicated to Tabernaemontaneae and their specific MIA metabolism. With more than 2,000 scaffolds, we first observed that the current assembly of the *T. elegans* genome is highly fragmented, as are those of *V. thouarsii* and *P. lamerei* (Cuello, Stander, Hans J Jansen, *et al*., 2022; Cuello *et al*., 2024).

These three species thus share a high percentage of transposable elements (TEs), which probably explains the largest genome size of Tabernaenemontaneae compared to Vinceae species. Such a correlation has already been proposed in plants and more generally in eukaryotes (Kidwell, 2002; Lee and Kim, 2014; Dai *et al*., 2022). As a matter of facts, we noted that assembling genomes with a high TE fraction can be challenging without scaffolding techniques such as chromatin contact capture (Hi-C) or optical mapping data. In contrast, assembling genome of Apocynaceae species with a low TE content - such as *C. roseus* (v2.1, 173 scaffolds), *V. minor* (v1, 296 scaffolds), and *R. tetraphylla* (v1, 76 scaffolds) - achieved intermediate scaffolding levels without the use of Hi-C (Cuello, Stander, Hans J. Jansen, *et al*., 2022; Stander *et al*., 2022; Stander *et al*., 2023). It is therefore highly likely that the high TE content explains the low level of scaffolding observed in our assembled *T. elegans* genome. However, despite the high fragmentation of the *T. elegans* assembled genome, the synteny between *C. roseus* and *T. elegans* confirms their close evolutionary relationship and demonstrates the robust conservation of genome organization (Figure 2C).

We have used this new genomic resource to gain insights into the metabolism of MIAs with a particular focus on pachysiphine biosynthesis. Indeed, pachysiphine and related MIAs, including the dimer conophylline, exhibit valuable pharmacological properties that would be difficult to exploit without the development of sustainable production strategies independent from the natural plant resources (Umezawa and Shirabe, 2021; Zhou *et al*., 2023; Courdavault *et al*., 2020). This highlights the necessity of elucidating the biogenetic modalities of these compounds starting with the C6-C7 epoxidation of tabersonine. Interestingly, a similar modification occurs in the synthesis of lochnericine albeit with the epoxide in the opposite orientation (Carqueijeiro, Brown, *et al*., 2018). On this basis, we mined the *T. elegans* genome for orthologues of TEX but none of the tested candidates displayed the expected pachysiphine synthase activity. By contrast, PYS1 and PYS2, two orthologues of T3O, which catalyzes the C2-C3 epoxidation of tabersonine en route to vindoline, synthesize pachysiphine as confirmed by NMR and by transient overexpression in *C. roseus* (Figure 3C-D, Figure 5J, Figure S7-10). Given the phylogenetic distance between the TEX-like and T3O-like enzymes (Figure S3), this suggests that they are two independent evolutionary scenarios for acquiring the capacity to epoxidize the C6-C7 double of tabersonine. It is thus highly likely that TEX evolved from an ancestor common with T16H through gene duplication and specification, whereas a similar phenomenon occurred for PYS1 and PYS2 from an ancestor with T3O common. Such a distinct origin for enzymes with very close activities is similar to the evolution of vincadifformine 19-hydroxylase (V19H) and T19H in *C. roseus*. Indeed, both enzymes hydroxylate the ethyl group of tabersonine and vincadifformine, which differ only in the C6-C7 double bond. However, V19H originated from a T3O homolog, whereas T19H evolved from a different subclade (Figure S3) (Giddings *et al*., 2011; Williams *et al*., 2019). Consistent with this hypothesis, microsynteny and phylogenetic analyses suggest that the *PYS* genes are derived from CYP71D1 and CYP71D2 in *T. elegans* (Figure 2D). Their speciation in Tabernaemontaneae probably explains the emergence of the pachysiphine synthase activity since pachysiphine derivatives appear to be specific to this clade (Kam *et al*., 2003). Above all, it also suggests that the T3O-like subclade of P450s should be of interest for the whole MIA metabolism and that new biosynthetic enzymes from this clade will be discovered soon. In addition, the syntenic region PYS2 hosts a gene encoding a putative NMT. Albeit this enzyme did not exhibit any *N*-methyltransferase activity on tabersonine and pachysipine (Figure 6A), it is highly possible that it N-methylates other MIAs given the high identity with the NMT identify in *C. roseus* (Figure S3C).

Interestingly, PYS1 and PYS2, although probably of different origins, have biochemical properties quite similar to those of TEX. While they readily accept tabersonine derivatives substituted on the indole ring, other modifications including C2-C3 epoxydation and 19-hydroxylation or 19-acetylation prevent PYS1 and PYS2 activity. Although these latter modifications do not occur in *T. elegans*, they may indicate that the catalytic sites of PYS1/PYS2 adopt a specific conformation that allows only limited substrate access. The modeling of PYS thus strongly suggests that the specific orientation of tabersonine in the substrate binding site could be determined by the position of a serine residue that is delocalised between PYS and T3O, while the narrow channel of PYS above the heme could restrict the substrate SOM. Interestingly, a careful examination of the models and sequences of other P450s involved in MIA biosynthesis, such as T19H, V19H, alstonine synthase (AS) and sarpagan bridge enzyme (SBE) revealed the presence of a conserved serine residue in all active sites (Dang *et al*., 2018; Williams and De Luca, 2023). Although in different positions, this serine may represent a critical residue for a controlled activity on MIA. In addition, several of these P450s (T16H, T19H, AS and SBE) also have bulk residues and a proline, potentially influencing the channel aperture, which controls access to the heme and thus the enzyme activity. Docking experiments also suggest that Pi-interactions and van der Waals forces are involved in the stabilization of tabersonine in the correct orientation in the binding sites of PYS and T3O. Identifying such residues thus enrich our knowledge of MIA enzymology as a whole but also opens up new perspectives for protein engineering and the development of enzymes with broader/adapted substrate specificities or improved activities as already reported for other MIA biosynthetic enzymes such as strictosidine synthase and strictosidine ß-D-glucosidase (Chen *et al*., 2006; Zhang *et al*., 2022).

Beyond leading to the identification of PYS1 and PYS2, getting access to the *T. elegans* genome should also greatly enhance our ability to identify new other MIA biosynthetic enzymes or isoforms. For instance, while there are no or only few biosynthetic gene clusters in this genome as well as in other MIA-producing plants, the careful examination of the genomic environment of *PYS* revealed the presence of a gene encoding a close homologue of NMT that may strengthen the hypothesis of its potential role IN MIA biosynthesis. In addition, screening the genome for *O*-methyltransferases based on the homology with *C. roseus* 16OMT gave access to three new enzymes, Te16OMT1, Te16OMT2 and Te16OMT3, which were active on both tabersonine and pachysiphine (Figure 6 and Figure S19). While the 16OMT activity is widespread in MIA-producing plants including *C. roseus* and *V. minor* (Stander *et al*., 2020; Levac *et al*., 2008), *T. elegans* is the only species described to have three active copies of this enzyme to date. The close proximity of the corresponding gene in the genome also strongly suggests that these enzymes arose recently from local gene duplication, as compared to *C. roseus,* which carries two distant copies (Lemos Cruz *et al*., 2023). This suggests that different evolutionary processes were involved in the demultiplication of 16OMTs in *C. roseus* and *T. elegans*. Such a difference in the evolution of the isoform copies also exists for T16H, which are present in two contiguous copies in *C. roseus* as a result of local duplication whilst only one TeT16H exists (Figure 3A).

Finally, the identification of PYS1 and PYS2 allowed the development of microbial cell factories producing pachysiphine derivatives (Figure 7). Based on the observed substrate acceptance of these enzymes (Figure 5I), we created two different yeast strains expressing T16H/16OMT/PYS and T19H/TAT, respectively. Their successive cultivation allowed the tailor-made production of 16-methoxy-19-acetylpachysiphine. To date, no plant has been reported to express a similar set of enzymes. For instance, *C. roseus* expresses T16H/16OMT and T19H/TAT/TEX in different organs, whereas *T. elegans* only co-expressesT16H/16OMT/PYS (Carqueijeiro, Brown, *et al*., 2018; Carqueijeiro, Dugé De Bernonville, *et al*., 2018). Thus, 16-methoxy-19-acetylpachysiphine s a new-to-nature MIA whose potential pharmacological properties deserve to be compared with those of pachysiphine. In addition to broadening the toolbox for MIA engineering, securing the bioproduction of pachysiphine and derivatives through metabolic engineering offers an alternative source for these MIAs. Given the high accumulation of tabersonine in *Voacanga africana* seeds, scaling up the production of pachysiphine and derivatives from tabersonine bioconversion (Kikura-Hanajiri *et al*., 2009; Koroch *et al*., 2009).

In conclusion, in this work we provide the genomic sequence of *T. elegans* that we used for the discovery of PYS. This identification highlights the different evolutionary processes that have been recruited to epoxidize tabersonine at the C6-C7 position, but in opposite orientations. Overall, this also enriches our understanding of the diversification and speciation of the MIA metabolism and opens new avenues for the MIA metabolic engineering.

## EXPERIMENTAL PROCEDURES

### Sample collection, DNA extraction and sequencing

*Tabernaemontana elegans* seeds obtained from Boutique végétale (https://boutique-vegetale.com/) were germinated and planted in individual pots. Seedlings were allowed to grow at 28°C under a 16h light/8h dark cycle. DNA was extracted from young leaves of two-month-old plants using Plant DNeasy kit (Qiagen) following the manufacturer’s instructions. DNAseq library was performed by Future Genomics Technologies (Leiden, The Netherlands) using Nextera Flex kit (Illumina) for Illumina sequencing and ONT 1D ligation sequencing kit (Oxford Nanopore Technologies Ltd) for Nanopore sequencing. Illumina libraries were sequenced in paired-end mode (2 × 150 bp) using Illumina NovaSeq 6000 technology. ONT libraries were sequenced on Nanopore GridION and PromethION flowcells (Oxford Nanopore Technologies Ltd) with the guppy version 4.2.3 high-accuracy basecaller. A total of 98,230,674 reads were obtained from the Illumina NovaSeq 6000 sequencing, 6,466,085 from the ONT PromethION sequencing and 142,157 from the ONT GridION sequencing.

### *De novo* genome assembly

Future Genomics Technologies (Leiden, The Netherlands) assembled the genome of *T. elegans*. ONT reads were assembled using Flye (v.2.8.3,Kolmogorov *et al*., 2019). Illumina reads were used twice to polish contig using pilon (v.1.23, Walker *et al*., 2014). Raw whole genome sequencing data and assembled genome have been deposited in the NCBI database under BioProject accession number XXXXXX.

### RNA extraction and sequencing

RNA was extracted from liquid nitrogen flash-frozen roots, young and old leaves, flowers and floral buds using NucleoSpin RNA Plant and Fungi mini kit (Macherey-Nagel) and purified using RNase-free TURBO DNase set (Thermo Fisher Scientific), both according to the suppliers’ instructions. RNA library construction and 150 bp paired-end sequencing was performed at FGTech using Illumina NovaSeq 6,000 technology. Raw RNA-seq data have been deposited in the NCBI database under BioProject accession number XXXXXX.

### Gene model prediction and functional annotation

RNA-seq reads were processed using FastP (v0.20, Chen *et al*., 2018) with default settings. The resulting reads were then aligned to the assembled *T. elegans* genome using HiSAT2 with -a -q –maxintron-len 5000 parameters (2.2.1, Kim *et al*., 2019). Sequence alignment map files (SAM) were sorted and converted to binary alignment files (BAM) to finally be merged and sampled with samtools (1.15.1, Danecek *et al*., 2021). Each individual alignment was then assembled into individual transcriptomes using StringTie with default parameters (v2.1.7, Pertea *et al*., 2015). The resulting individual transcriptomes were then merged into a non-redundant set of 49,533 transcripts related to 25,045 protein coding genes using stringtie --merge. Coding sequences were then identified within transcripts using Transdecoder (v5.7.1).

### Assembly quality assessment

The quality of *T. elegans* genome assembly was evaluated by MerQury (v.1.7, Rhie *et al*., 2020) and the LTR assembly index (LAI) from LTR-retriever (v.2.9.6, Ou and Jiang, 2018). The completeness of the genome assembly and annotation was assessed with Benchmarking Universal Single-Copy Orthologs (BUSCO v.5.4.2, Simão *et al*., 2015) using default setting and a plant-specific database containing 2,326 orthologs (eudicots_odb10). Statistics for the gene models were obtained using the seqIO library from biopython (v.1.83).

### Transposable element annotation and whole genome duplication analysis

The sensitive mode of the extensive *de novo* transposable elements (TE) annotator (EDTA v.1.9.5, Ou *et al*., 2022) was used to identify and annotate TE. Whole genome duplication (WGD) events were assumed using the DupPipe pipeline (Barker *et al*., 2010).

### Orthology analysis and phylogenetic tree reconstruction

*T. elegans*, *V. thouarsii* (Cuello, Stander, Hans J Jansen, *et al*., 2022), *R. tetraphylla* (Lezin *et al*., 2023), *C. roseus* (Li *et al*., 2023), *V. minor* (Stander *et al*., 2022), *P. lamerei* (Cuello *et al*., 2024), *G. sempervirens* (Franke *et al*., 2019), *O. pumila* (Rai *et al*., 2021), *S. lycopersicum* (Hosmani *et al*., 2019), *C. acuminata* (Kang *et al*., 2021), *V. vinifera* (Shi *et al*., 2023) and *A. thaliana* (Lamesch *et al*., 2012) proteomes were filtered to conserve proteins of at least 30 amino acids. Proteomes were further subjected to CD-HIT (v.4.7, Fu *et al*., 2012) and the longest representative protein was retained in each CD-HIT cluster. The resulting protein datasets were inputting into OrthoFinder (v.2.5.4, Emms and Kelly, 2019) using the following parameters: -S diamond -M msa -A muscle. An approximate maximum-likelihood phylogenetic tree was then inferred from 488 single-copy orthogroups. We then analyzed orthogroup expansion and contraction across the phylogenetic tree using Cafe5 (v.4.2.1, Mendes *et al*., 2021).

### Genome colinearity analysis

MCscan pipeline (Tang, Wang, *et al*., 2008; Tang, Bowers, *et al*., 2008) together with the JCVI utility library (Tang *et al*., 2024) was used to detect synteny blocks. Circos (v0.69-8) was used to highlight synteny links.

### Candidate gene cloning

Genomic DNA was extracted from flash-frozen young leaves using NucleoSpin Plant II kit (Macherey-Nagel) as per manufacturer’s instructions. The full length sequences of the candidate genes were amplified from genomic DNA using nested PCR : (i) firstly, candidate genes were amplified with 5’ and 3’ untranslated region specific primers of each (Table S6) and (ii) these PCR products were used as a template for a PCR with gene specific primers (Table S6). PCR were done by using the Q5 High-Fidelity DNA Polymerase (New England Biolabs) and PCR products were purified by the NucleoSpin PCR clean-up kit (Macherey-Nagel). Golden gate reactions were performed to clone the 33 candidate genes and GFP into the pHREAC vector (Addgene; plasmid #134908) or a modified pHREAC containing BsmBI site instead of BsaI, by using BsaI-HF-v2 or BsmBI-v2 golden gate assembly kit (New England Biolabs). For golden gate reactions, 50 ng of pHREAC and insert were added with 10 units of BsaI-HF-v2 or BsmBI-v2 golden gate mix and the T4 DNA ligase buffer at 1X into 10 μl of reaction using these parameters: [(2 min - 37°C; 5 min - 16°C) x35; 5 min - 60°C, 5 min - 80°C]. Cloning reactions were transformed into thermocompetent *Escherichia coli* TOP10 and recombinant colonies were selected on LB agar plates with kanamycin (50 μg.mL^−1^). To select positive clones, colony PCRs were performed by using the DreamTaq DNA polymerase (Thermo-Scientific) and sequencing primers designed into the pHREAC vector (Table S6). Positive colonies were inoculated into 5 ml of LB medium with kanamycin (50 μg.mL^−1^) for overnight cultures at 30°C in order to isolate cloned plasmids using the Nucleospin plasmid kit (Macherey-Nagel). Identities of the cloned sequences were confirmed by Sanger sequencing using pHREAC sequencing primers (Table S6). Then, recombinant vectors were used to transform electrocompetent GV3101 *Agrobacterium tumefaciens* cells by electroporation. Recombinant cells were selected on LB-agar plates with kanamycin (50 μg.mL^−1^), rifampicin (50 μg.mL^−1^) and gentamicin (20 μg/ml).

### Transient expression of candidate genes in *Nicotiana benthamiana*

Recombinant *A. tumefaciens* were grown in 3 ml LB medium cultures complemented with kanamycin (50 μg.mL^−1^), rifampicin (50 μg.mL^−1^) and gentamicin (20 μg.mL^−1^) at 28°C for 24 h. LB cultures of 15 ml without antibiotic were inoculated at 1/250 from the pre-culture and let grown 16 h at 28°C. Cells were harvested by centrifugation for 10 min at 2,000 g, washed with 5 ml of infiltration buffer (MES 10 mM pH 5,6; MgCl2 10 mM) and resuspended in 5 ml of infiltration buffer. For enzymatic assays of T3O-like, TEX-like or NMT-like candidate genes, *Agrobacterium* solutions were diluted to OD600 of 0,3 in 10 ml of infiltration buffer supplemented with acetosyringone at 150 μM and further incubated 3 h at 28°C before infiltrations. For enzymatic assays of 16OMT-like candidate genes, *Agrobacterium* solutions of OMT candidates were independently diluted to an OD600 of 0,3 in infiltration buffer supplemented with an *Agrobacterium* culture of TeT16H at an OD600 of 0,3 and acetosyringone at 150 μM and further incubated as described above. For substrate specificity of PYS1, *Agrobacterium* solutions of PYS1, T3O, T19H and/or TAT were cultured or co-cultured to an OD600 of 0,3 as mentioned above. Infiltrations were performed using 1 ml syringes into the leaf abaxial side of 4 week-old *N. benthamiana* younger leaves.

### Enzymatic assays and PYS1 substrate specificity with *N. benthamiana* disks

Four days post-infiltration, five leaf disks of 1 cm diameter were collected in the younger infiltrated leaf of each plant. These disks were added into a well of a 24 well plate with 500 μl of infiltration buffer and tabersonine at 100 μM. The solution was infiltrated into leaf disks with a vacuum chamber at 50 mBar. The 24 well plate was incubated at 26°C with 16 h of light during 24 h. Infiltration medium was collected and diluted at 1/5 in methanol 100%, clarified by centrifugation (15 min, 18,000 g, 4°C) and analyzed by liquid chromatography combined to mass spectrometry.

### Enzymatic production of pachysiphine, purification and NMR analyzes

The *Agrobacterium* strain transformed with pHREAC-PSY1 (Tel01G.15088) was used to agro-infiltrate 50 plants of *N. benthamiana* like the transient expression for enzymatic assays. Four days after infiltration, 4 leaves per plant were harvested and cut in 1 cm squares. Leaf pieces were added in 600 ml of infiltration buffer containing 20.18 mg of tabersonine (100 µM). The solution was infiltrated into leaf pieces with a vacuum chamber at 50 mBar and incubated at 26°C with 16 h of light during 24 h on a shaking plate. After incubation, leaf pieces were discarded and the infiltration buffer containing enzymatic products was collected and centrifuged for 15 min at 17,000 g. An alkaloid liquid/liquid extraction protocol was applied to this resulting buffer (600 ml) to extract pachysiphine. The buffer (200 ml) was alkalinized to pH10 with NaOH solution at 1M and extracted four times with 75 ml of ethyl acetate. This was repeated twice for the remaining buffer. After drying the organic phase, the extract (56 mg) was resuspended in MeOH (23 mg.mL^−1^) and purified by preparative HPLC (Interchim, puriFlash 5.250), with a C18 column (Xselect HSS T3 OBD, 5µM, 10 x 250 mm). A 90-minute method at a flow rate of 4 mL.min^−1^ was applied including a 60-minute gradient from 20 % B (A: H_2_O, B: MeCN) to 60 % B, then a 30-minute gradient from 60% B to 95% of B. Collected fractions were analyzed by UPLC-MS, assembled and dried to obtain 5.2 mg of pure pachysiphine.

Nuclear magnetic resonance (NMR) analyzes were performed at 300 MHz for ^1^H and 75 MHz for ^13^C with BRUKER AVANCE 300 spectrometer. Chemical shifts are given in parts per million (δ) relative to the solvent peak (respectively 7.26 and 77.00 for ^1^H and ^13^C in CDCl_3_).

^1^H NMR (CDCl_3_, 300 MHz) : δ 8.98 (br, 1H), 7.18-7.12 (m, 2H), 6.90-6.83 (m, 1H), 6.81 (d, *J* = 7.7 Hz, 1H), 3.78 (s, 3H), 3.59 (dd, *J* = 12.9, 1.1 Hz, 1H), 3.26 (d, *J* = 3.8 Hz, 1H), 3.07 (bd, *J* = 3.8 Hz, 1H), 3.05-2.98 (m, 1H), 2.89 (d, *J* = 12.9 Hz, 1H), 2.75-2.67 (m, 2H), 1.39 (d, *J* = 15.5 Hz, 1H), 2.46 (d, *J* = 1.6 Hz, 1H), 2.09 (dd, *J* = 11.5, 6.5 Hz, 1H), 1.70 (dd, *J* = 11.5, 4.2 Hz, 1H), 1.02-0.90 (m, 2H), 0.73 (t, *J* = 7.2 Hz, 3H). ^13^C NMR (CDCl_3_, 75 MHz) : δ 169.0, 165.3, 143.1, 137.7, 127.8, 121.5, 120.5, 109.4, 91.2, 71.2, 56.3, 54.6, 52.2, 51.12, 51.06, 49.7, 47.1, 37.2, 26.4, 23.3, 7.2.

### Transient overexpression of PYS1 in *Catharanthus roseus*

*Catharanthus roseus* plants were infiltrated in triplicates with *Agrobacterium* strains containing either *PYS1* or *GFP* genes as described by Koudounas *et al*., 2022. Five days post-infiltration, the apex part of younger infiltrated leaves were harvested, weighted and flash frozen in liquid nitrogen. Then, methanol 0.1% of formic acid was added into samples and they were sonicated 45 min to extract metabolites. Supernatants were separated from the residual powder by centrifugation and diluted in methanol to analyze by LC-MS.

### Ultra-Performance Liquid Chromatography combined to Mass Spectrometry analyzes

UPLC-MS analysis were performed with an ACQUITY UPLC (Waters) system coupled with a SQD2 mass spectrometer (Waters), operating with an electrospray ionization source. All the system was monitored by Masslynx 4.2 software (Waters). Compounds were separated into a Waters Acquity HSST3 C18 column (150 x 2.1 mm, 1.8 μm) with an injection volume of 5 μl into the column setting at 55°C and 0.4 mL.min^−1^ of solvent flow rate. The mobile phase was composed of solvent A (0.1% formic acid in water) and solvent B (0.1% formic acid in acetonitrile), following an 18 min linear gradient from 10% to 50% of solvent B. Mass spectrometer detections were performed in a positive mode with a protonated mass to charge ratio (*m/z*) ([molecular mass + H]+). The different MIAs analyzed on LC-MS were described with their *m/z* ratios and their retention times (Table S7).

UPLC-HRMS/MS analyzes were carried out on an AQUITY UPLC system (Waters) coupled to a quadrupole time-of-flight mass spectrometer (Synapt G2-Si HDMS, Waters) equipped with an electrospray ionization interface (ESI). Chromatographic separations were performed on a Waters Acquity BEH C_18_ (100 x 2.1 mm, 1.7 µm) column, and the temperature was maintained at 40°C. Mixtures of H_2_O (A) and ACN (B) with 0.1% formic acid were eluted at a flow rate of 0.4 mL.min^−1^ with a gradient of 10% to 60% of (B) in 12 min. The samples (1 µL injection volume) were analyzed in fast data-dependent acquisition (fDDA) mode consisting of a full MS survey scan from 50 to 1200 *m/z* (scan time = 0.2 ms) followed by MS/MS scans for the three most intense ions (*m/z* 100-1200; scan time = 0.05 ms). A collision energy ramp was set from 10 to 40 eV for low-masses and 40 to 90 eV for high-masses. Acquisition and data processing were achieved using MassLynx version 4.2. Files were transformed into mzML format with MSConvert software, part of the ProteoWizard package (Chambers *et al*., 2012). Comparison figures of two MS/MS spectra were processed with R version 4.4.1 (R Core Team, 2023) with the package Spectra (Rainer *et al*., 2022).

#### Homology modeling and protein-ligand docking prediction

Three-dimensional models of PYS1, PYS2 and T3O were generated by homology-based modeling on TrRosetta using templates (Du *et al*., 2021). Models with the highest TM-score were assessed for prediction quality with MolProbity (Williams *et al*., 2018), where hydrogens were also added to the proteins. The catalytic heme was manually positioned in the catalytic site of the enzymes by superimposing the models with 5ylw and 7cb9 crystal structures, which were the best hits used as templates during the modeling process. Docking studies for tabersonine were conducted using Autodock Tools and Vina (Eberhardt *et al*., 2021) within a grid box of 20 x 20 x 20 Å centered on the capacity above iron heme and using exhaustiveness of 8. The predictions showing tabersonine SOM (C2-C3 for T3O and C6-C7 for PYS) within 5 Å of the heme iron, were retained among the top ten poses ranked with calculated binding affinity. Control experiments were performed with an homology based model of CYP76AH1 from *Sativa miltiorrhiza* to evaluate the specificity of tabersonine poses in PYS1, PYS2 and T3O. Protein-ligand interactions were visualized with PyMOL.

#### Yeast strains construction

The coding sequence of CrCPR2, T16H1, 16OMT, T19H, TAT and PYS1 were amplified with Phusion High-Fidelity DNA polymerase (Fermentas) and specific primers (Table S6). The PCR products were either cloned by conventional digestion/ligation procedure in the SpeI or NheI cloning site (Fermentas) or by Golden Gate assembly (New England Biolabs) of pDPTA125, pDPTA111, pDPAC124, pPETA104, pPETA102 and pPEAC115 donor vectors under constitutive promoters as indicated in Table S8 (Kulagina *et al*., 2022). *Saccharomyces cerevisiae* CEN.PK113-7D (MATa MAL2-8C, SUC2) was used as the starting strain (Van Dijken *et al*., 2000). CRISPR-Cas9 integration method was employed to introduce expression cassettes in yeast hot-spot locus (Mikkelsen *et al*., 2012) as described in Kulagina *et al*., (2022). Briefly, yeast strains harboring the pCfB2312 vector were co-transformed with gRNA helper plasmids and the appropriate NotI-linearized donor DNA vectors, by using the lithium acetate transformation method (Chen *et al*., 1992). Yeast transformants were selected on YPD medium plates (20 g.L^−1^ of peptone, 10 g.L^−1^ of yeast extract, and 20 g.L^−1^ of glucose) supplemented with G418 (200 μg.mL^−1^) and nourseothricin (100 μg.mL^−1^), and screened for the effective integration of expression cassette by colony PCR. The yeast strains constructed and used in this study are listed in Table S9.

#### Production of new-to-nature tabersonine derivatives in yeast

The “TOP” strain pre-culture overnight in 5 mL YPD was diluted to OD600 of 1 in 200 µL of YPD supplemented with tabersonine at 40 µM. The “TOP” culture was incubated at 30°C under constant agitation at 200 rpm during 48 h. Every 24 h and 48 h of culture, 22 µL of glucose 20% (w/v) and YP 10X were added to the yeast cultures. Enzymatic products released by yeasts into the culture medium were centrifuged at 3,000 g during 5 min in order to be used for a second bioconversion step. Then, a “TT” strain pre-culture overnight in 5 mL YPD was started and diluted to OD600 of 1 in 200 µL of YPD. These 200 µL were centrifuged at 3,000 g during 5 min to remove the supernatant. This “TT” strain pellet was solubilized into the kept “TOP” supernatant and incubated at 30°C under constant agitation at 200 rpm during additional 24 h. Cultures were centrifuged at 17,000 g during 15 min to remove cells. Supernatant was diluted at ⅕ in methanol and analyzed by UPLC-MS.

#### Tests of PYS1 substrate specificity with tabersonine derivatives in yeast

To test the substrate specificity of the PYS1 with tabersonine, 16-hydroxytabersonine and 16-methoxytabersonine, the “PYS1” strain pre-culture overnight in 5 mL YPD was diluted to an optic density at 600 nm of 1 in 200 µL of YPD supplemented with standards of these 3 substrates at 40 µM. The “PYS1” culture was incubated at 30°C under constant agitation at 200 rpm during 48 h. Enzymatic products were released by yeasts into the culture medium so the culture was centrifuged at 17,000 g during 15 min to remove cells. Supernatant was diluted at ⅕ in methanol and analyzed by UPLC-MS. To test the substrate specificity of the PYS1 with 2,3-epoxytabersonine, 19-hydroxytabersonine and 19-acetyltabersonine, the 3 yeast strains named “T3O”, “T19H” and “TT’’ were pre-cultured overnight in 5 mL of YPD. Pre-cultures were independently diluted to an optic density at 600 nm of 1 in 200 µL of YPD supplemented with tabersonine at 40 µM. The 3 strain cultures were incubated at 30°C under constant agitation at 200 rpm during 24 h. Cultures were centrifuged at 3,000 g during 5 min to keep supernatants and enzymatic products. Then, a “PYS1” strain pre-culture overnight in 5 mL YPD was started and diluted to an optic density at 600 nm of 1 in 200 µL of YPD. These 200 µL were centrifuged at 3,000 g during 5 min to remove the supernatant. This “PYS1” strain pellet was solubilized into the kept “T3O”, “T19H” and “TT” supernatants and incubated at 30°C under constant agitation at 200 rpm during 48 h with addition of glucose 20% and YP 10X as described before. Cultures were centrifuged at 17,000 g during 15 min to remove cells. Supernatant was diluted at ⅕ in methanol and analyzed by UPLC-MS as described before.

#### Subcellular localization studies

To analyze the subcellular localizations of PYS1 and PYS2 in *N. benthamiana*, these two P450 genes were cloned in fusion with the yellow fluorescent protein (YFP) on the protein C-terminal part into the pHREAC. The cloning procedure was already explained above in the “candidate gene cloning” part, except that specific primers described in Table S6 were used to amplify the *PYS1/2* and the *YFP* genes in order to form fusion proteins. Another plant expression vector was used with a gene that encodes a marker of the endoplasmic reticulum, named *CD3-953*, in fusion with the cyan fluorescent protein (*CFP*) gene as described by Nelson *et al*., (2007). These three recombinant vectors were electroporated in *A. tumefaciens* GV3101 as described above to produce three different strains named ER-CD3-953-CFP, PYS1-YFP and PYS2-YFP. Each strain were prepared as mentioned above but before agro-infiltration, the ER-CD3-953-CFP strain was co-cultured with an OD600 nm of 0,2 in the infiltration buffer with either PYS1-YFP or PYS2-YFP at an OD600 nm of 0,2. The leaf abaxial side of 4 week-old *N. benthamiana* younger leaves were agro-infiltrated to obtain transient overexpressions of the ER-CD3-953-CFP marker with PYS1-YFP or PYS2-YFP. At three days post-infiltration, infiltrated leaves were used to prepare microscope slides to observe the leaf abaxial side. Subcellular localizations were determined using an Olympus FV3000 confocal microscope. CFP fusion proteins were excited at 445 nm and emission was captured at 460-500 nm. YFP fusion proteins were excited at 514 nm and emission was captured at 530-630 nm. Chlorophyll was excited at 445 nm and emission was captured at 600-680 nm. The pattern of localization presented in this work is representative of circa 50 observed cells.

## ACCESSION NUMBERS

Assembled genome and related data have been deposited in the NCBI database under BioProject accession number XXXXXX. GeneBank access numbers are: PYS1, XXXXXX; PYS2, XXXXXX; TeT16H, XXXXXX; TeT16OM1, XXXXXX; TeT16OM2, XXXXXX, TeT16OM3, XXXXXX.

## ACKNOWLEDGEMENTS

This work was supported by EU Horizon 2020 research and innovation program [MIAMi project-Grant agreement N°814645]; APR IR program of the Région Centre-Val de Loire [ScaleBio project]; and ANR [project MIACYC – ANR-20-CE43-0010]. We are thankful to the chemical ecology platform of the Institut de Recherche sur la Biologie de l’Insecte (IRBI; Tours, France) for access to the UPLC/quadrupole time of flight mass spectrometer. We also thank Ascelin Choisy and Firdaous Fizazi for helping in gene cloning and pachysiphine purification, respectively.

## AUTHOR CONTRIBUTIONS

EL performed the cloning and testing of all candidate genes, protein subcellular localization and microbial cell factory development. MD and CC performed genomic analysis and homology searches. CBW and FM performed metabolic analysis and pachysiphine purification. ALLV, KK and AO contributed to candidate gene characterization. TP and NG performed the bioconversion experiment for the synthesis of new-to-nature MIAs. JP performed the NMR analysis. HJJ and RPD sequenced and assembled the *T. elegans* genome. SS, PLPA and MAB contributed to pachysiphine purification and NMR analysis. BSt-P and NGG contributed to subcellular localization studies. CS contributed to genome analysis. NP, MKJ and SOC contributed to experiment design and data analyses. SB designed experiments, performed data analyses and molecular modeling. VC conceived this study, designed experiments, contributed to data analysis and acquired funds. EL, MD, CC, SB and VC wrote the original manuscript. All other authors critically reviewed the manuscript.

## DATA AVAILABILITY STATEMENT

All data supporting the findings of this study are available within the paper and within its Supporting information published online. The genome annotation, its functional annotation, transcripts sequences, predicted CDS and protein sequences are available on the figshare XXXXXXXX.

## TEMPORARY ACCESS TO DATA

The genome annotation, its functional annotation, transcripts sequences, predicted CDS and protein sequences are temporarily available on the link below XXXXXXXXXX. Assembled genome and related data have been deposited in the NCBI database under BioProject accession number XXXXXXX and are temporarily available on the link below XXXXXXXXX.

## SUPPORTING INFORMATION

### List of supplemental tables

**Table S1**: Genome assembly features of *Tabernaemonta elegans*.

**Table S2**: Proportion of the different TE classes and family reported in the *T. elegans* genome.

**Table S3**: The 147 synteny blocks resulting from self synteny analysis of *T. elegans* genome.

**Table S4**: The 406 synteny blocks resulting from genome-wide synteny analysis between *T. elegans* and *C. roseus*.

**Table S5**: List of the putative MIA-related proteins identified by blastp.

**Table S6**: List of primers used in this study.

**Table S7**: List of analyzed MIAs on LC-MS

**Table S8:** List of temple-donor vectors used in this study.

**Table S9:** Yeast strains constructed and used in this study.

**Table S10**: List of protein sequences used to perform neighbor-joining phylogenetic trees.

### List of supplemental figures

**Figure S1:**
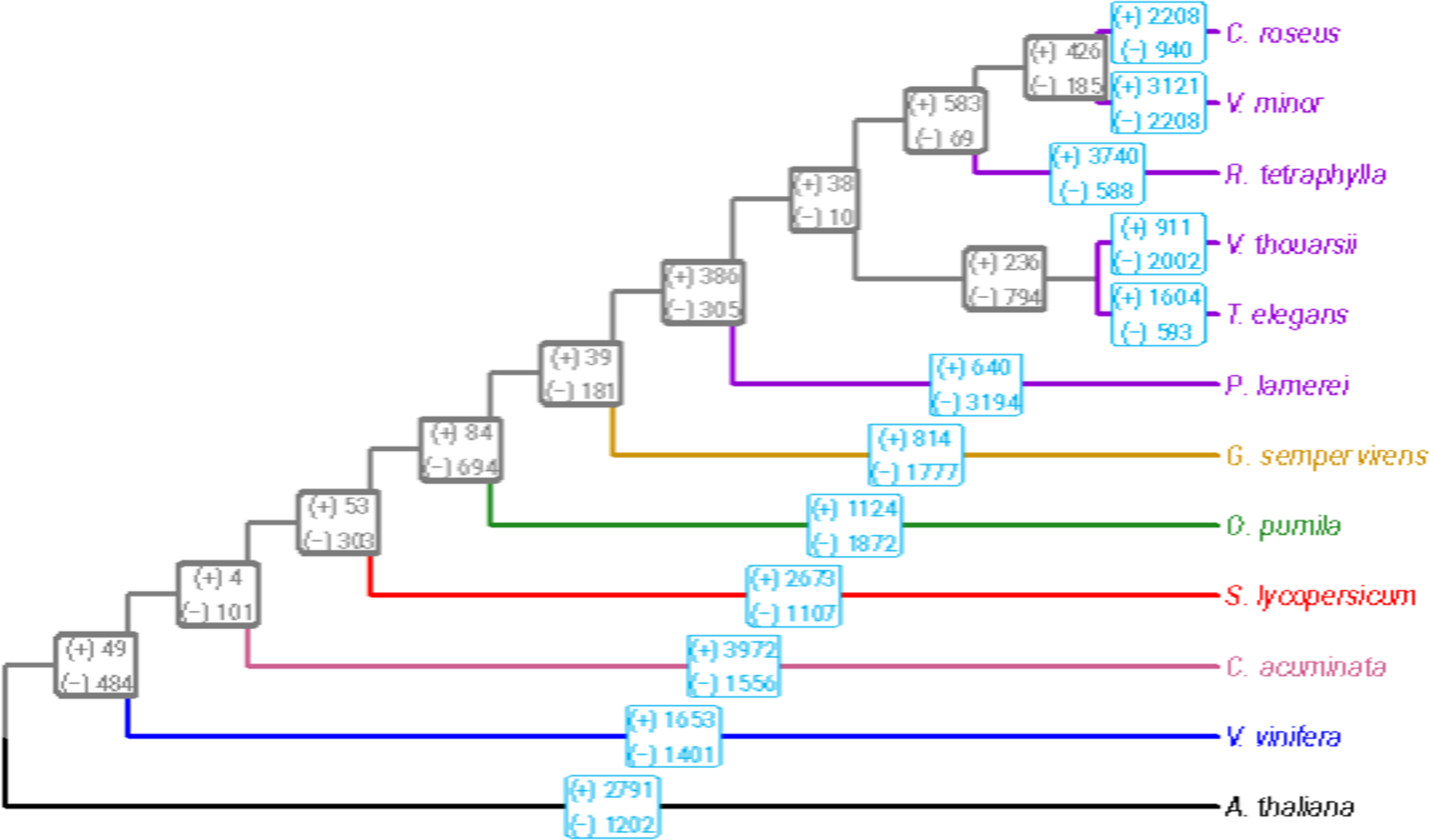
Phylogenetic tree of *T. elegans* and eleven other species, including five Apocynaceae (purple: *C. roseus*, *V. minor*, *R. tetraphylla*, *V. thouarsii* and *P. lamerei)*, one Gelsemiaceae (yellow: *G. sempervirens*), one Rubiaceae (green: *O. pumila*), one Solanaceae (red: *S. lycopersicum*), one Nyssaceae (pink: *C. acuminata*), one Vitaceae (blue: *V. vinifera*) and one Brassicaceae (black: *A. thaliana*). Gene family expansion (+) and contraction (−) were calculated using Cafe5 in each lineage (light bordered blue boxes) and in internal nodes of ancestral population for each taxon (thick bordered grey boxes).

**Figure S2:**
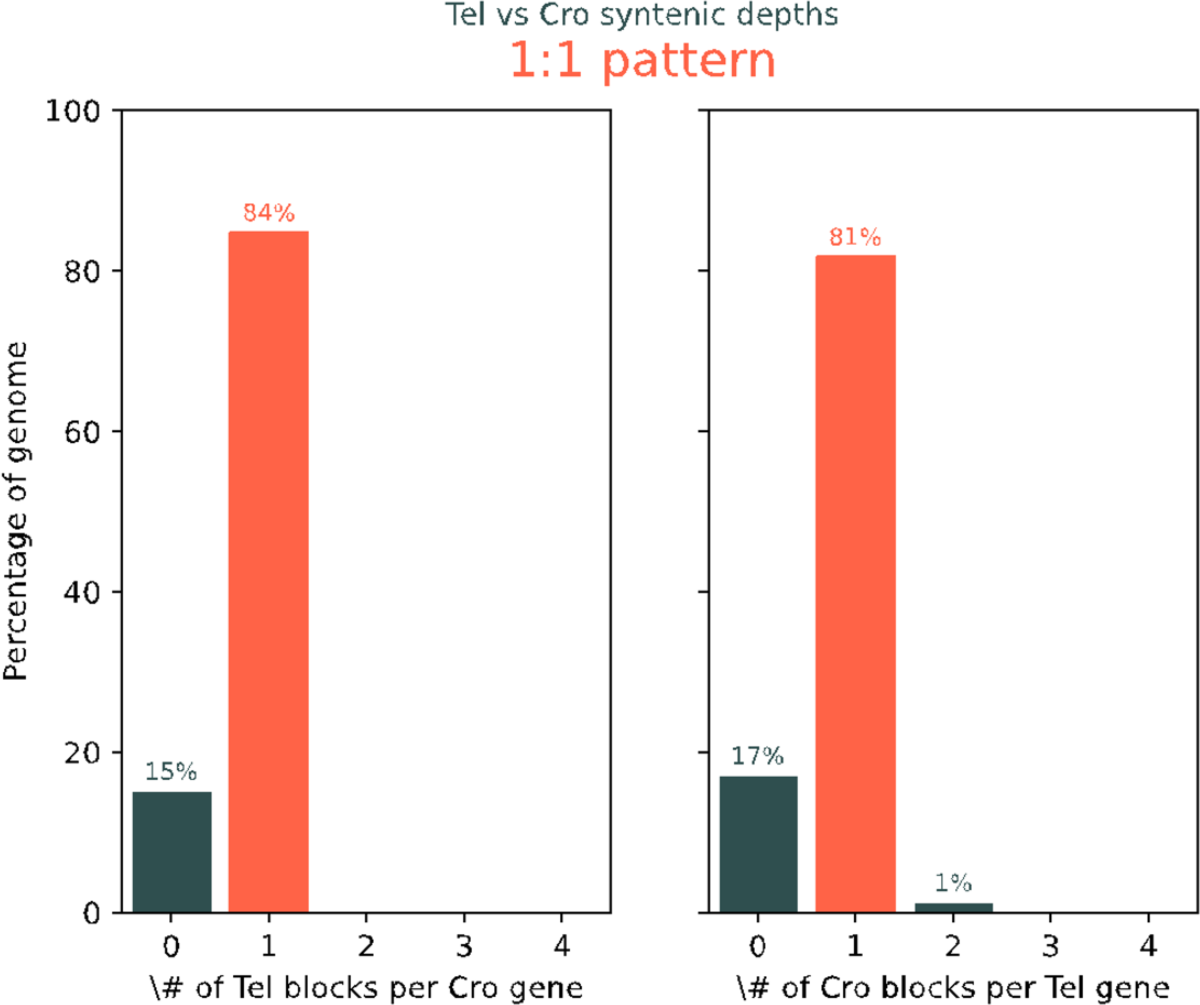
Syntenic pattern between *T. elegans* and *C. roseus* illustrating the 1:1 syntenic ratio. Left plot highlights proportion of reported number of *T. elegans* syntenic blocks per *C. roseus* gene. Right plot highlights proportion of reported number of *C. roseus* syntenic block per *T. elegans* gene.

**Figure S3:**
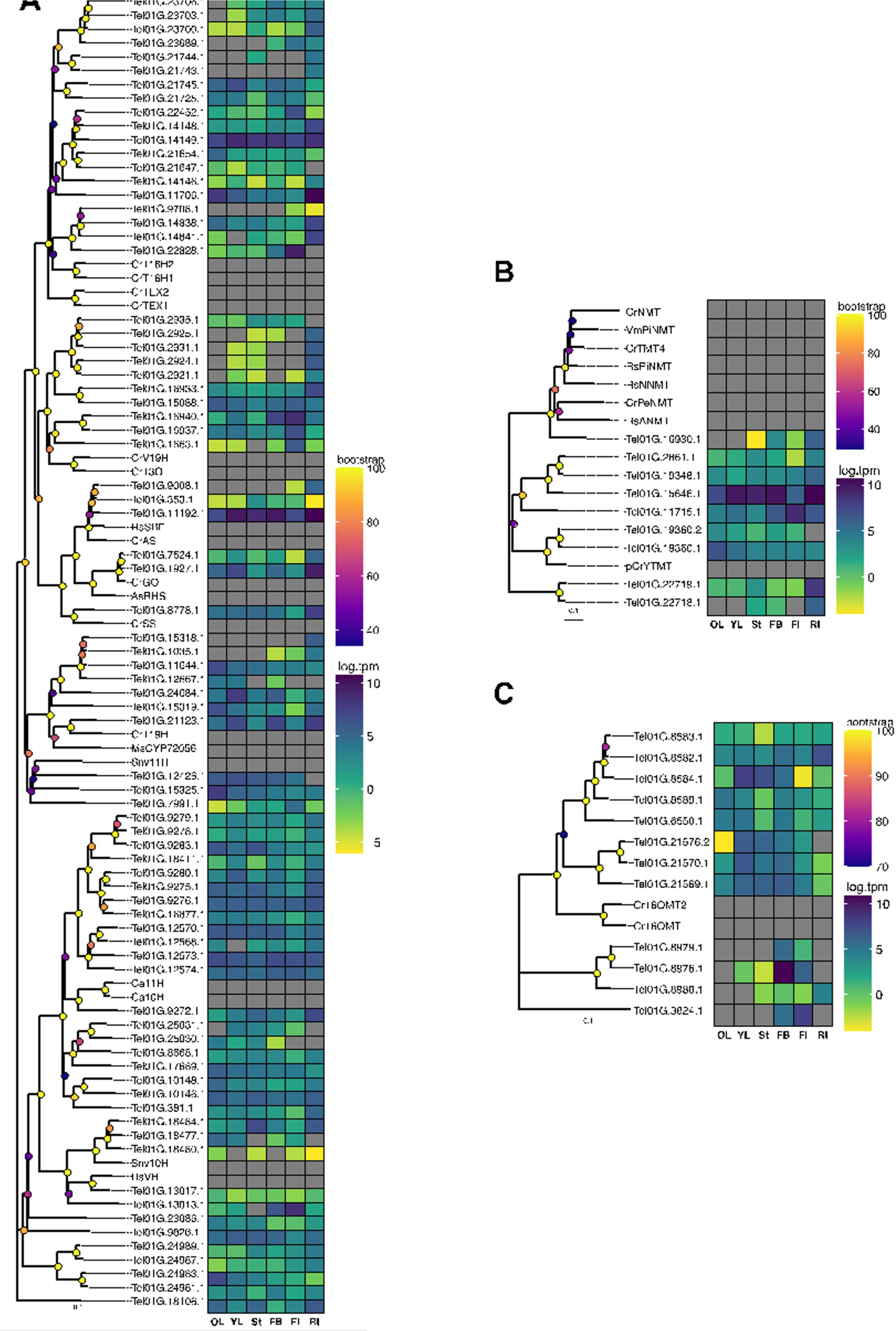
Gene expression in diverse plant tissues and phylogenetic analysis of TeCYP71s, TeOMTs and TeNMTs. (**A**) TeCYP71s, (**B**) TeOMTs and (**C**) TeNMTs, along with other reported enzymes involved in the biosynthesis of MIAs from diverse species, were categorized using a neighbor-joining phylogenetic tree (100 bootstrap replications). Metadata associated with bootstrap nodes is represented by a circle whose fill color scales proportionally with the bootstrap value while branch lengths scales proportionally with substitutions per site value. The scale bar represents 0.1 substitutions per site. Yellow to blue color gradient in heatmap represents gene expression displayed as log2(tpm) where yellow color represents lower expression and blue color higher expression.

**Figure S4:**
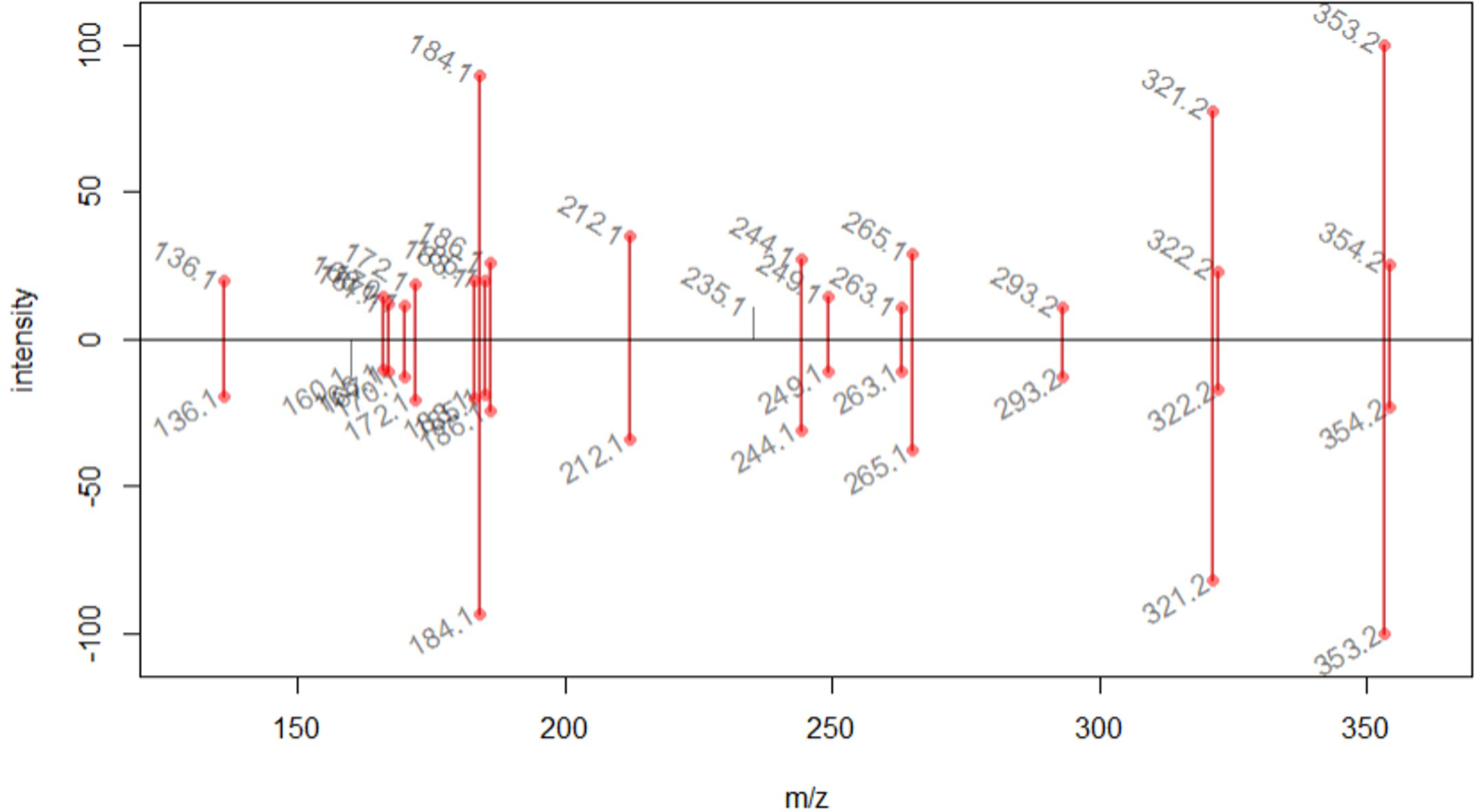
MS/MS comparison of authentic standard of 16-hydroxytabersonine [M+H]^+^ 353.1866 (top) and TeT16H product at *m/z* 353.1874 (bottom). Correlation score (GNPS-like score) of 0.97.

**Figure S5:**
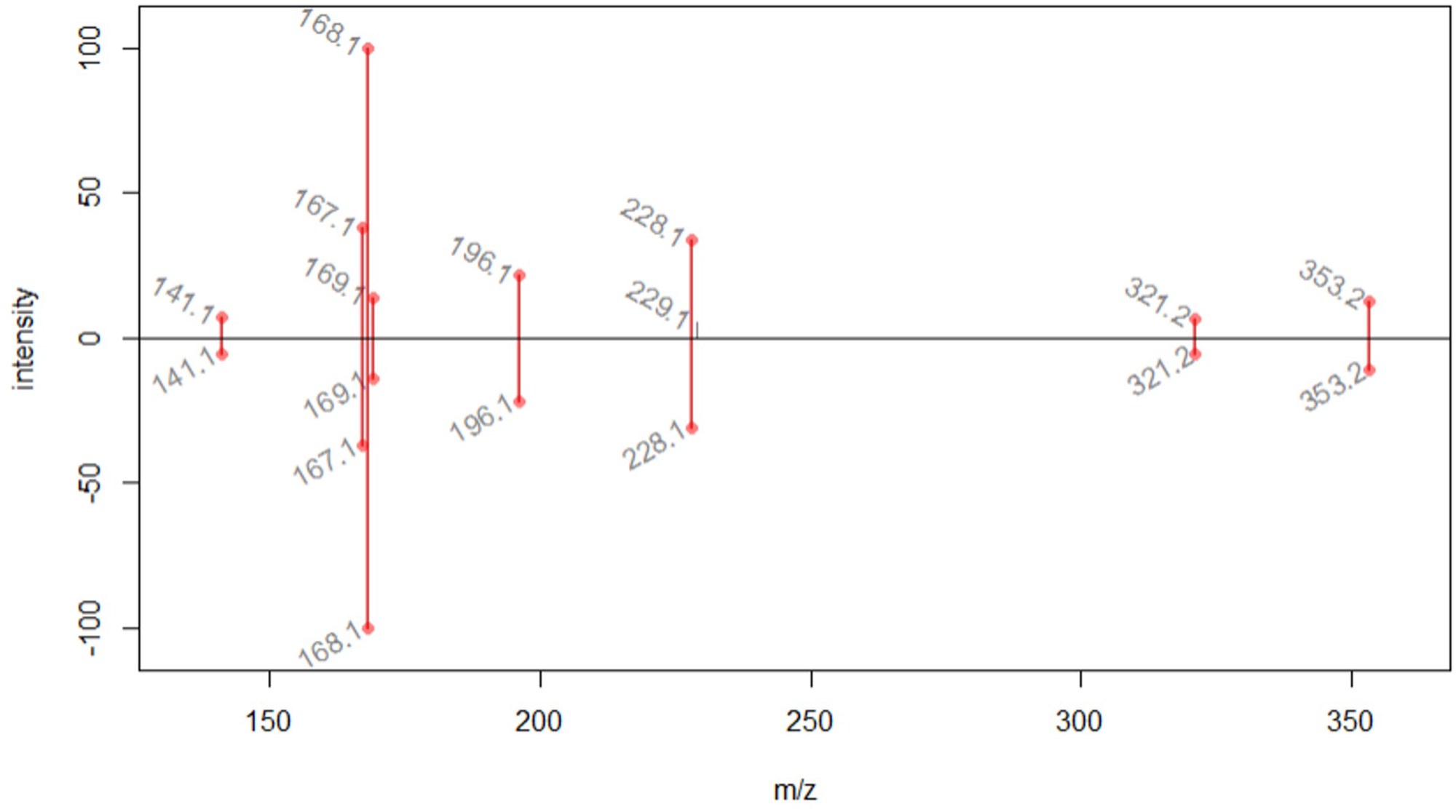
MS/MS comparison of authentic standard of pachysiphine [M+H]^+^ 353.1863 (top) and PSY1 product at *m/z* 353.1866 (bottom). Correlation score (GNPS-like score) of 0.98.

**Supplemental Figure S6:**
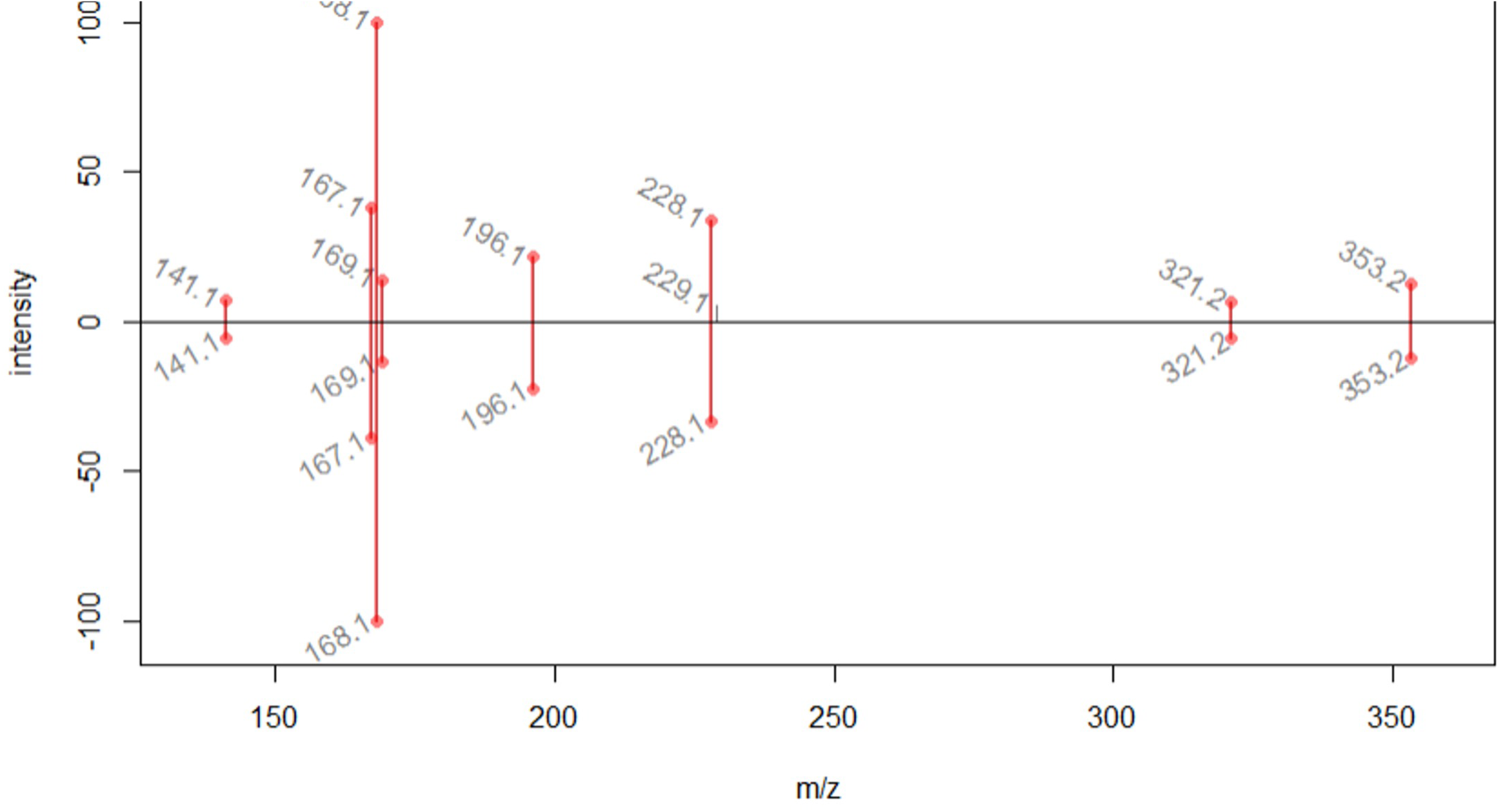
MS/MS comparison of authentic standard of pachysiphine [M+H]^+^ 353.1863 (top) and PSY2 product at *m/z* 353.1874 (bottom). Correlation score (GNPS-like score) of 0.98.

**Figure S7:**
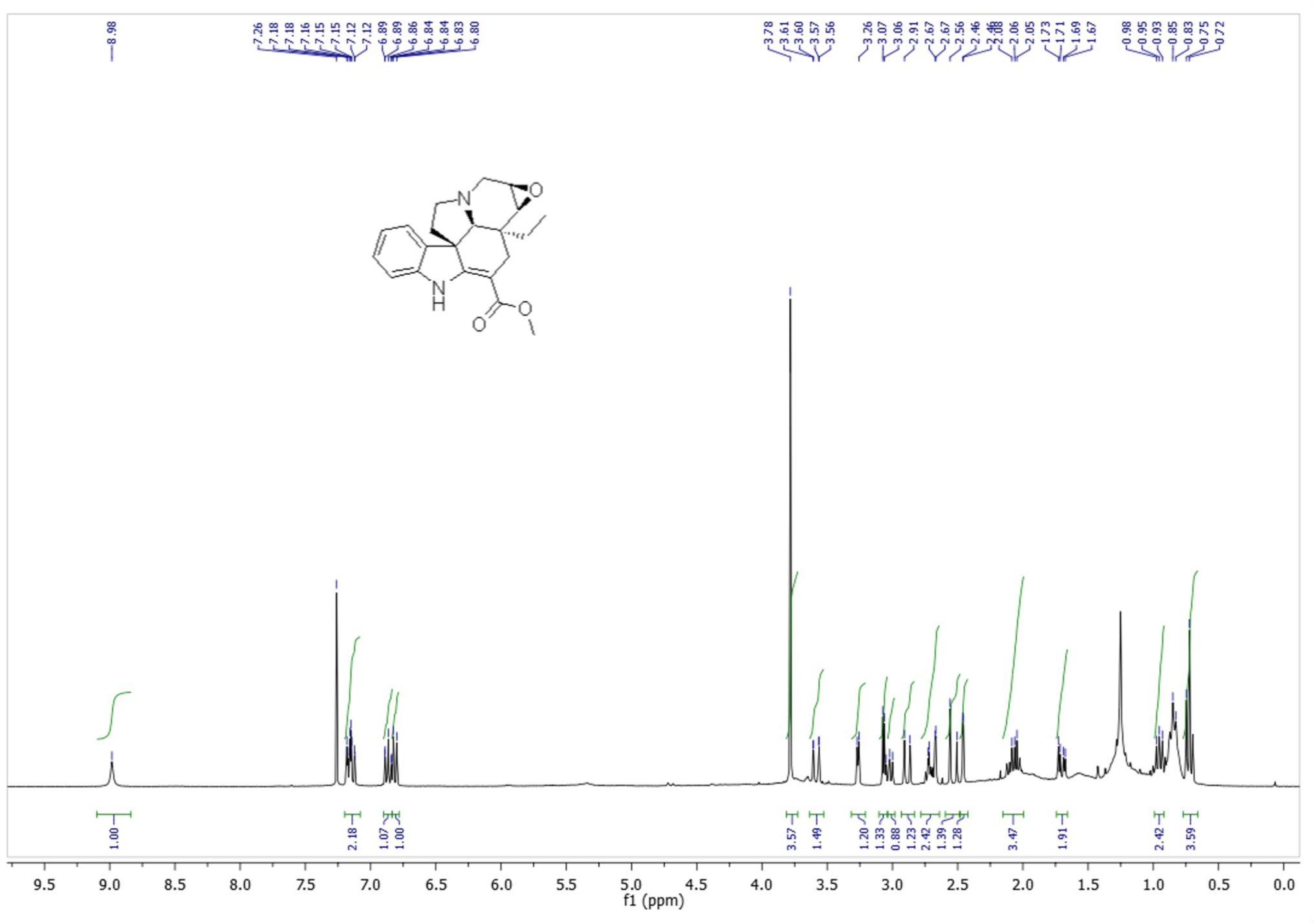
^1^H NMR (300 MHz) spectrum of (−)-pachysiphine in CDCl_3_

**Figure S8:**
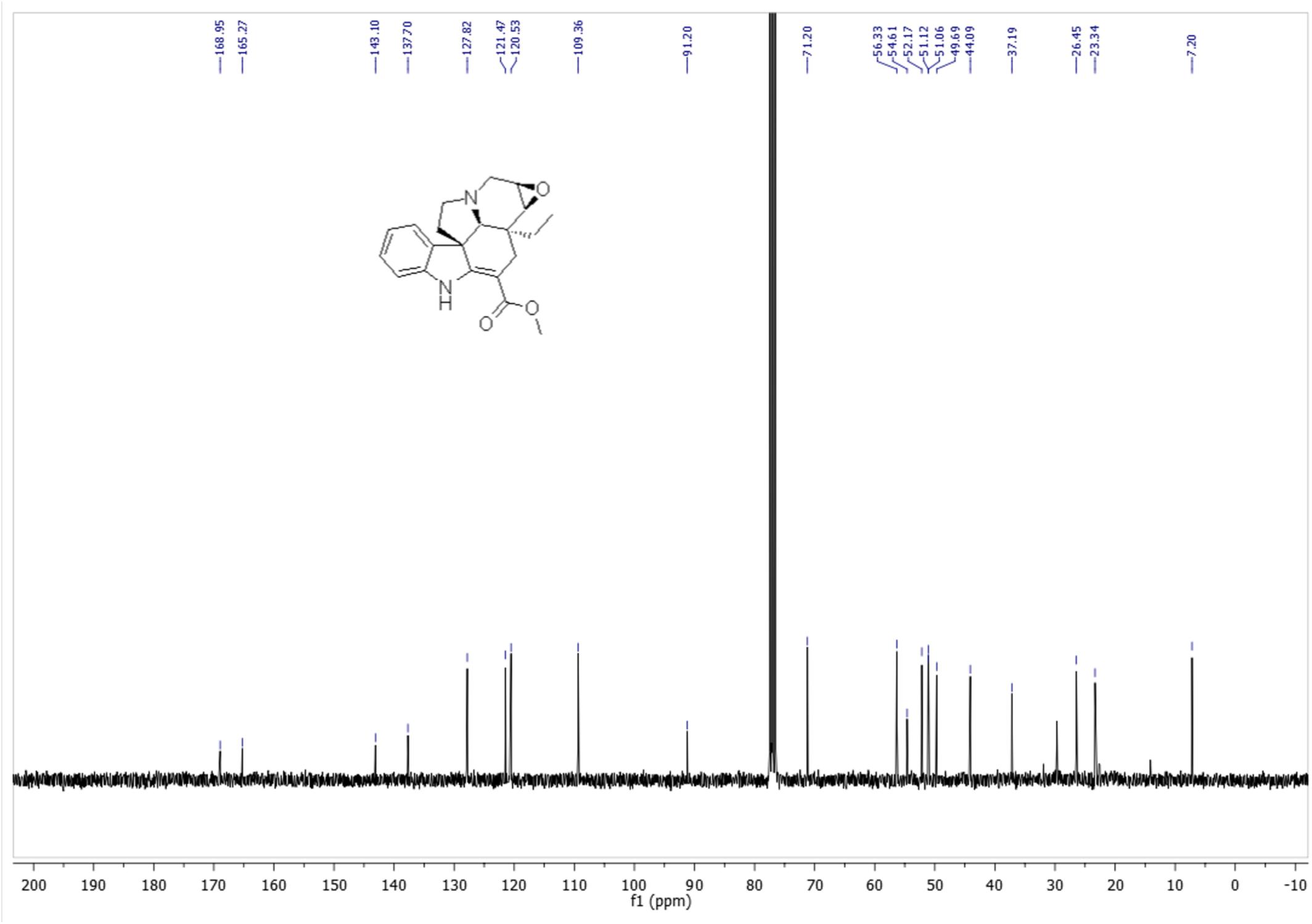
^13^C NMR (75 MHz) spectrum of (−)-pachysiphine in CDCl_3_

**Figure S9:**
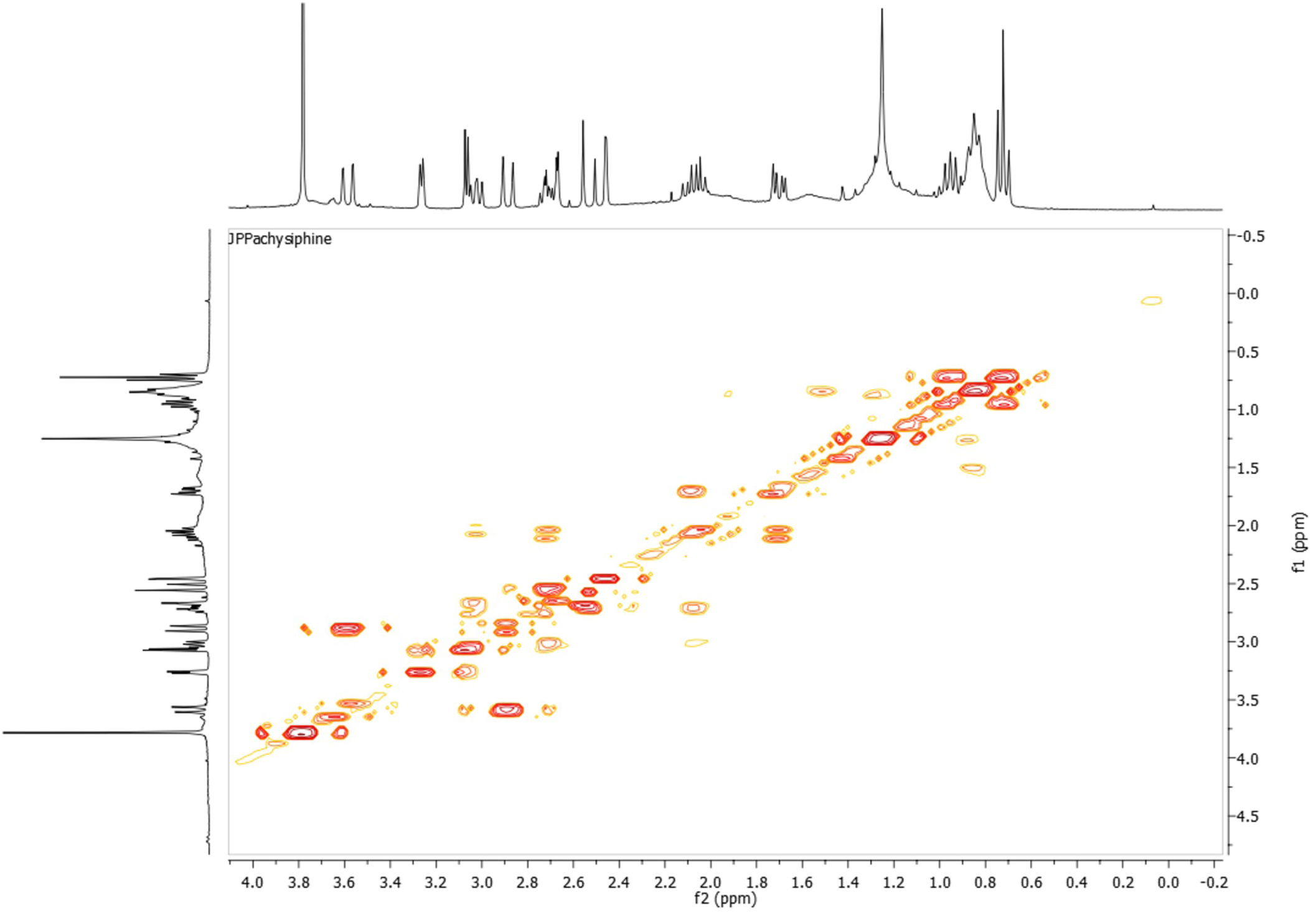
^1^H-^1^H COSY spectrum of (−)-pachysiphine in CDCl_3_

**Supplemental Figure S10:**
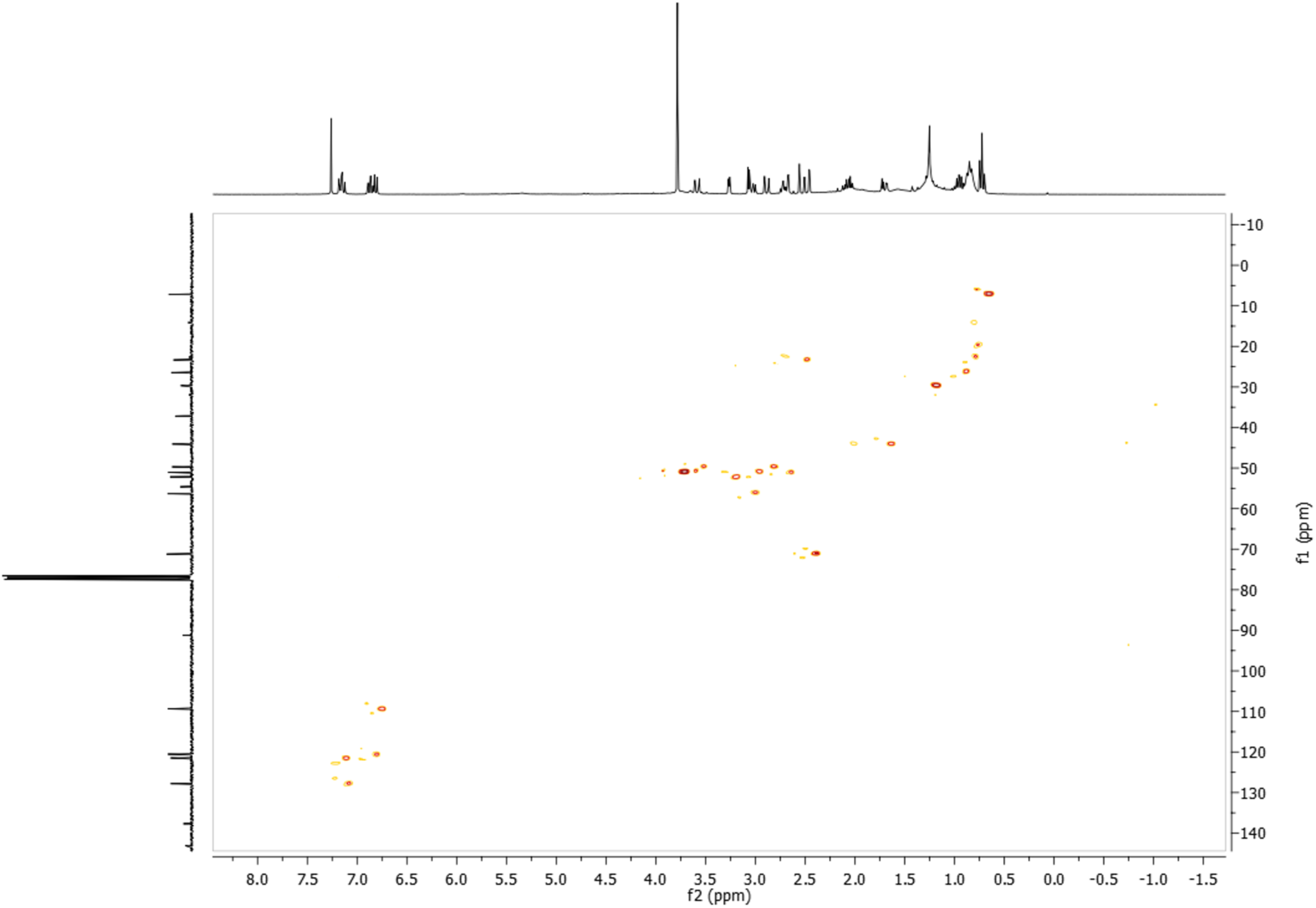
^1^H-^13^C HSQC spectrum of (−)-pachysiphine in CDCl_3_

**Figure S11:**
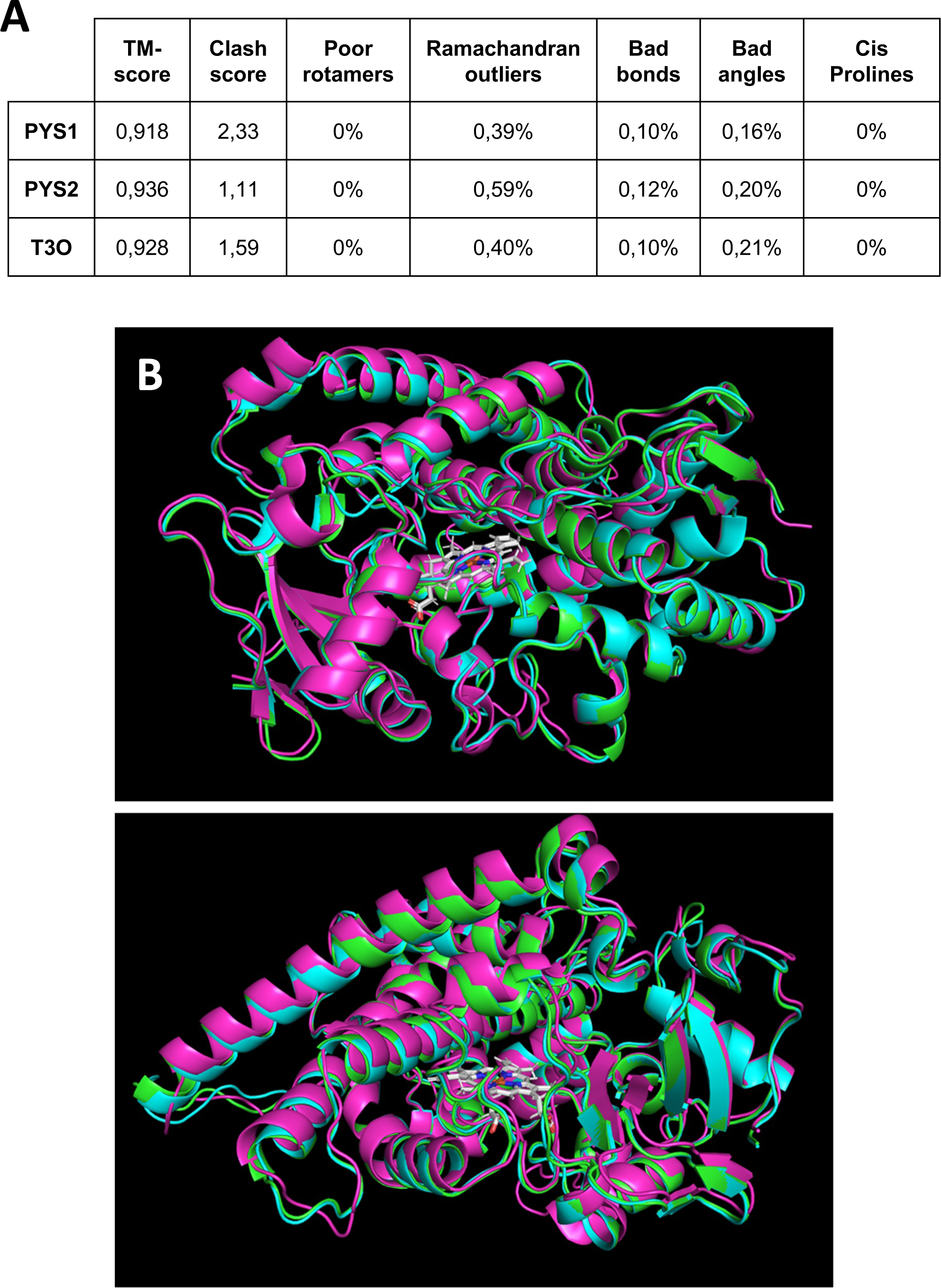
PSY1, PSY2 and T3O best homology-based three dimensional structure prediction from TrRosetta. (A) Qualitative assessments of models. (B) Superimposition of models with catalytic heme (white). PYS1 (green), PYS2 (blue) and T3O (pink). The transmembrane domain was removed as low confidence prediction parts of the proteins.

**Figure S12:**
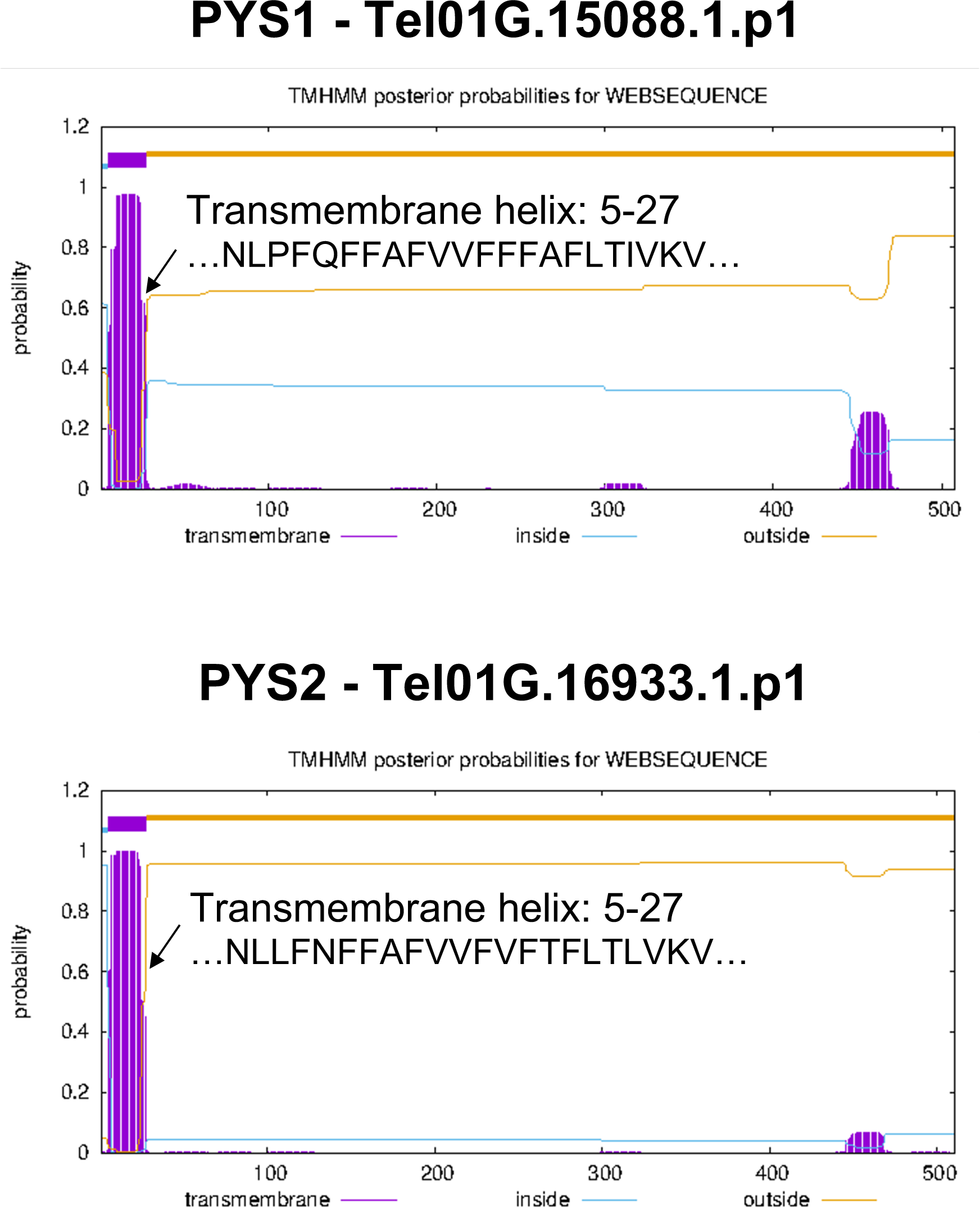
Detection of a putative transmembrane helix at the N-terminal end of PYS1 and PYS2. The probability of a residue to belong to a transmembrane helix has been calculated with a Markov model by the TMHMM server (https://services.healthtech.dtu.dk/services/TMHMM-2.0/).

**Figure S13:**
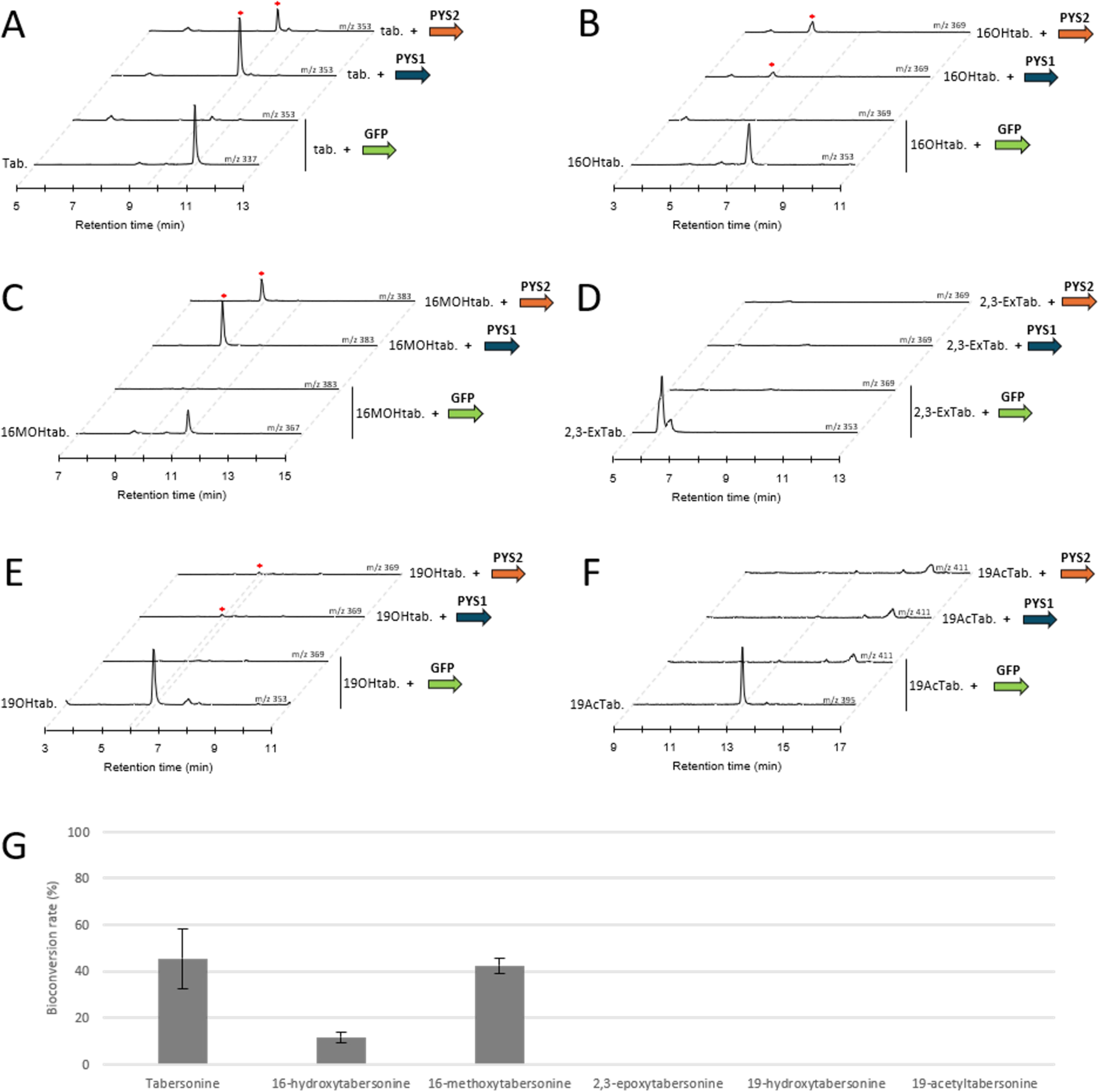
Substrate specificity of PYS1 and PYS2 with tabersonine derivatives in tobacco and yeast. (A-F) Substrate specificity of PYS1 and PYS2 in *N. benthamiana*. *PYS1* and *PYS2* genes or *GFP* were transiently overexpressed in tobacco leaves in presence of tabersonine derivatives and reaction products were analyzed by LC-MS. Positive differential mass chromatograms of oxygenated substrates are compared to GFP (negative control). Red asterisks indicate the formed products. (A) Enzymatic reactions with tabersonine (*m/z* 337), named “tab.”. (B) Enzymatic reactions with 16-hydroxytabersonine (*m/z* 353), named “16OHtab.”. (C) Enzymatic reactions with 16-methoxytabersonine (*m/z* 367), named “16MOHtab.”. (D) Enzymatic reactions with 2,3-epoxytabersonine (*m/z* 353), named “2,3-ExTab.”. (E) Enzymatic reactions with 19-hydroxytabersonine (*m/z* 353), named “19OHtab.”. (F) Enzymatic reactions with 19-acetyltabersonine (*m/z* 395), named “AcTab.”. (G) Substrate specificity of PYS1 in yeast with the same substrates mentioned above. Assays were done in triplicates to calculate the bioconversion rate after pick integrations.

**Figure S14:**
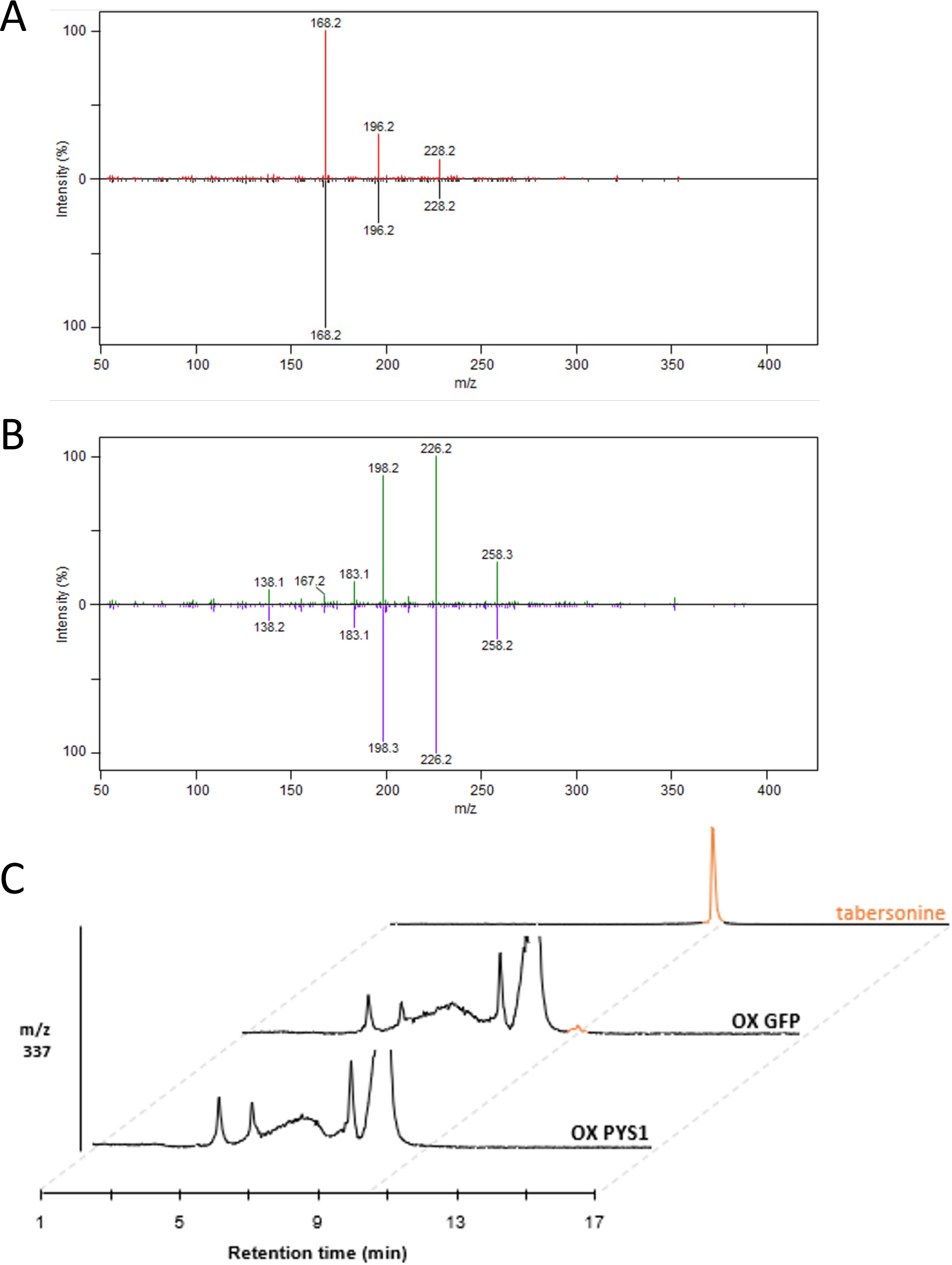
Overexpression of PYS1 in *Catharanthus roseus* leaves. *PYS1* or *GFP* genes were transiently overexpressed in *Catharanthus roseus* young leaves to analyze their MIA metabolisms by LC-MS. (A) MS/MS comparison of the molecular ion (*m/z* 353) of an authentic standard of pachysiphine (top) and the product at *m/z* 353 with the same retention time observed in *Catharanthus overexpressing PYS1* (bottom). (B) MS/MS comparison of the enzymatic product 16-methoxypachysiphine at m/z 383 obtained in yeast (top) and the product at *m/z* 353 with the same retention time observed in *Catharanthus overexpressing PYS1* (bottom). (C) In addition to the figure 5 J-L, positive differential mass chromatograms were analyzed at *m/z* 337 corresponding to the expected [M+H]^+^ of tabersonine standard (orange color) in *Catharanthus roseus* leaves overexpressing PYS1 or GFP. Tabersonine was barely detectable. *PYS: pachysiphine synthase. OX: overexpression. m/z: mass to charge ratio. LC-MS: liquid chromatography coupled to mass spectrometry*.

**Figure S15:**
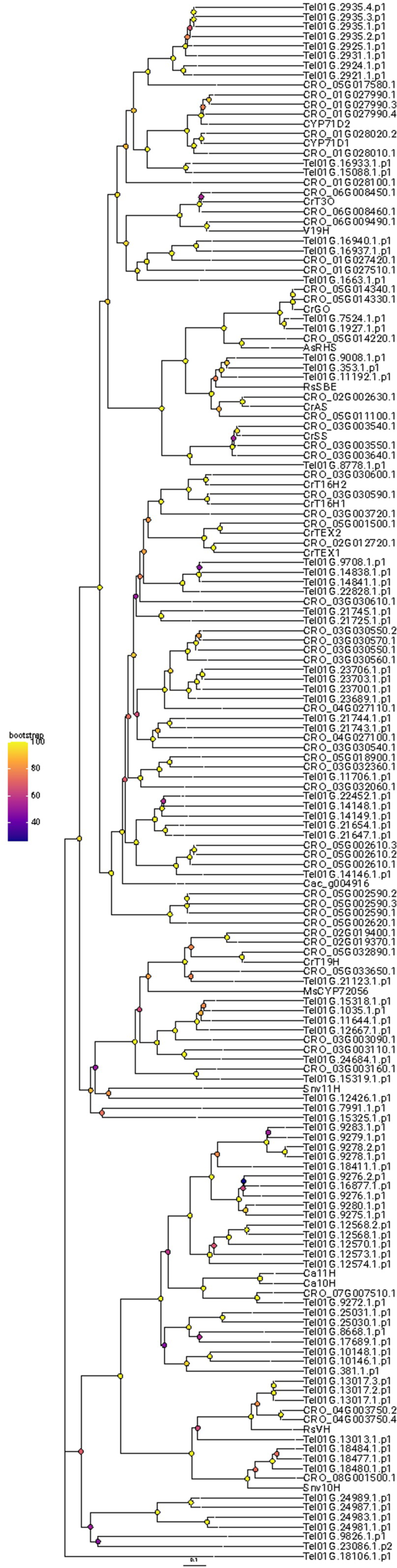
Phylogenetic analysis of TeCYP71s and CrCYP71s along with other reported CYP71 enzymes involved in the biosynthesis of MIAs from diverse species, were categorized using a neighbor-joining phylogenetic tree (100 bootstrap replications). Metadata associated with bootstrap nodes is represented by a circle whose fill color scales proportionally with the bootstrap value while branch lengths scales proportionally with substitutions per site value. The scale bar represents 0.1 substitutions per site.

**Figure S16:**
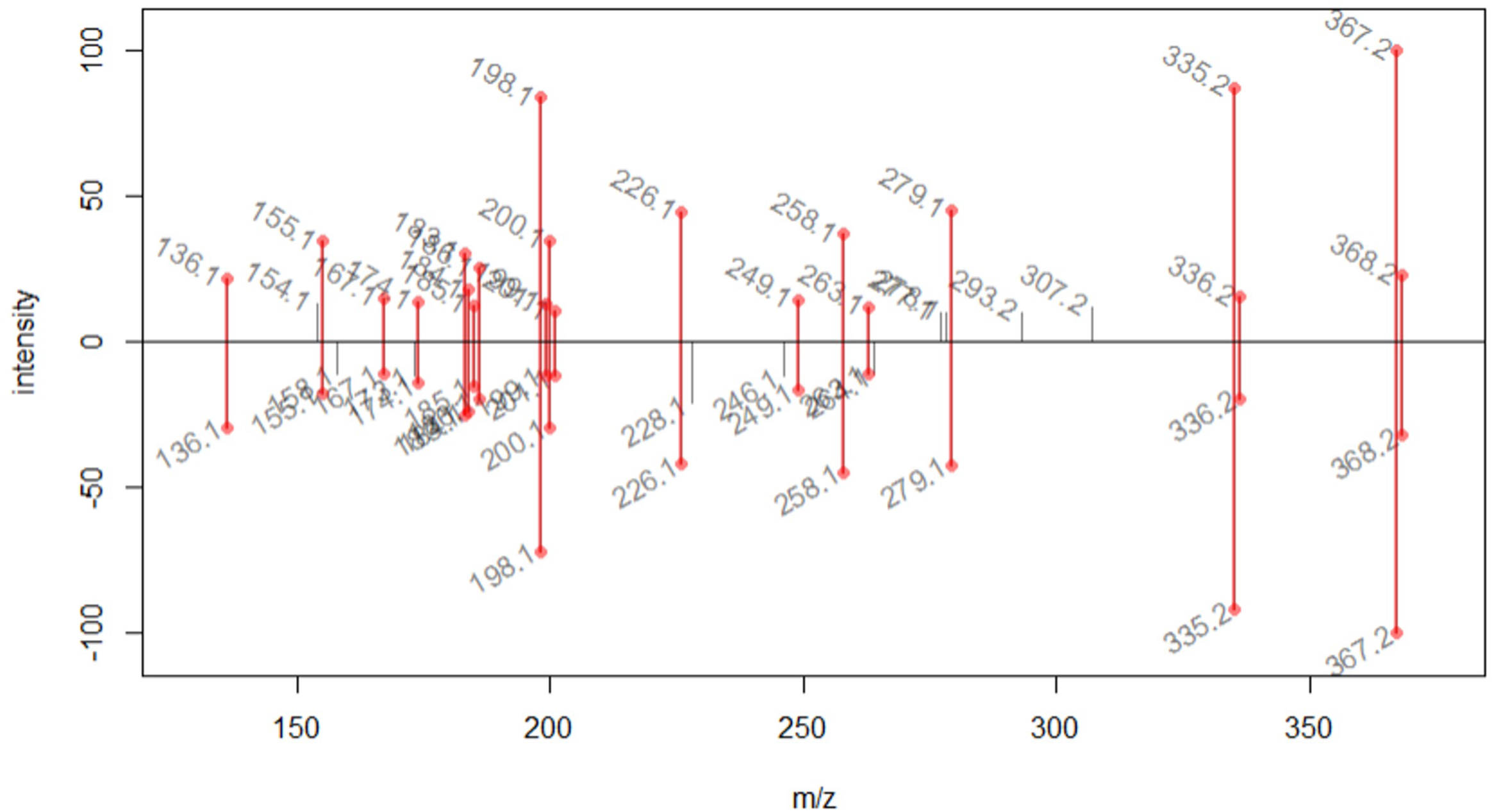
MS/MS comparison of authentic standard of 16-methoxytabersonine [M+H]^+^ 367.2021 (top) and TeT16H + Te16OMT1 product at *m/z* 367.2023 (bottom). Correlation score (GNPS-like score) of 0.91.

**Figure S17:**
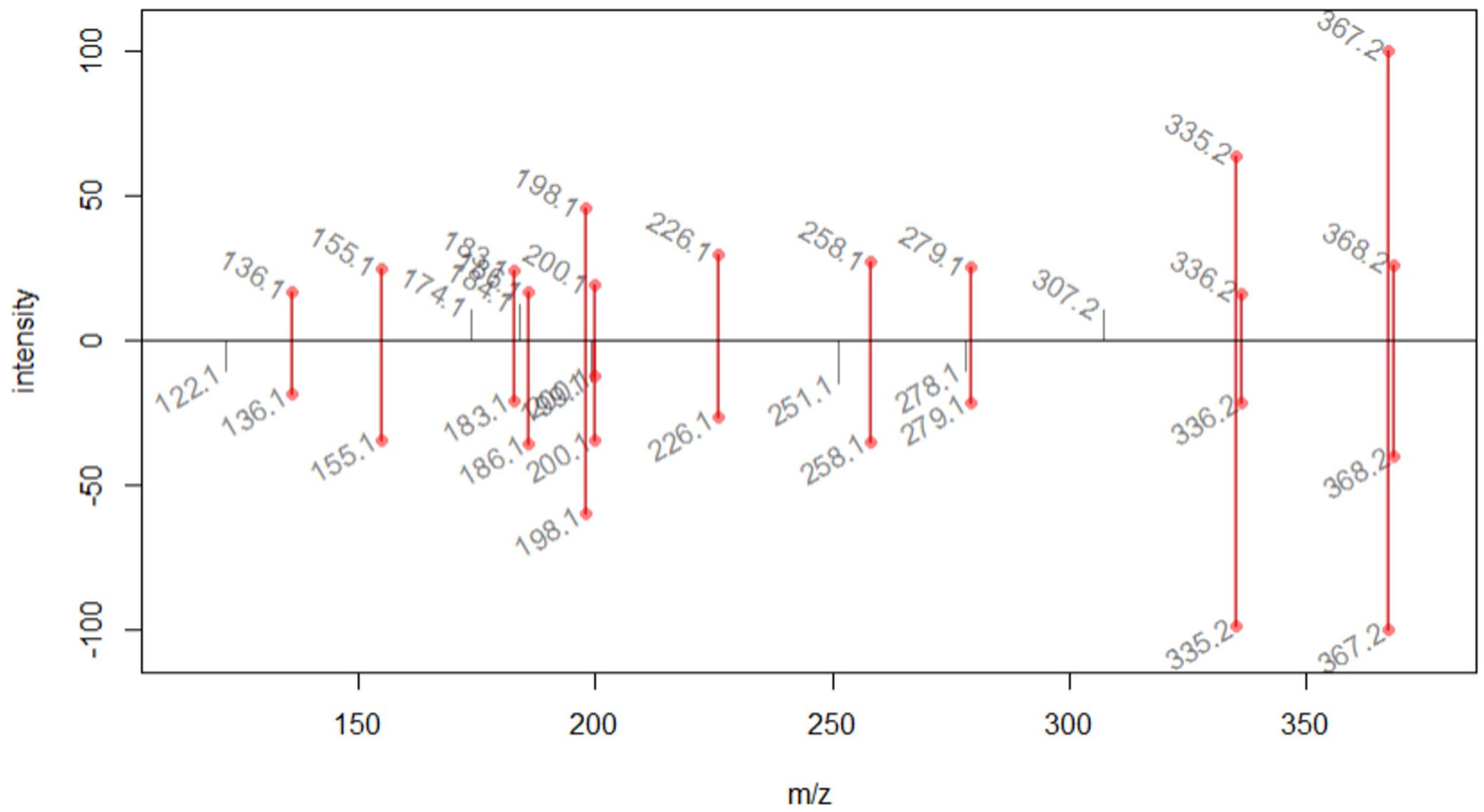
MS/MS comparison of authentic standard of 16-methoxytabersonine [M+H]^+^ 367.2021 (top) and TeT16H + Te16OMT2 product at *m/z* 367.2025 (bottom). Correlation score (GNPS-like score) of 0.99.

**Figure S18:**
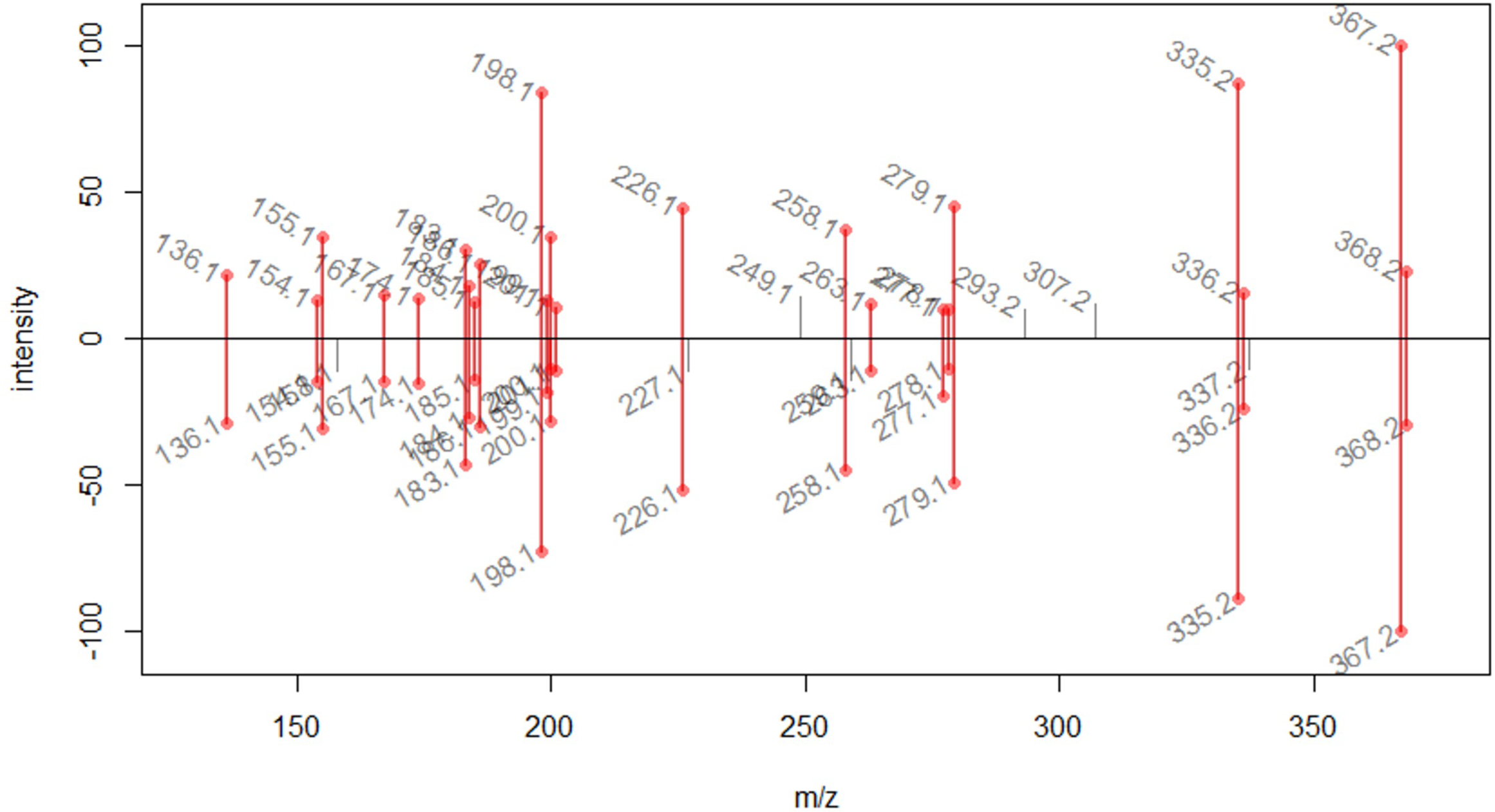
MS/MS comparison of authentic standard of 16-methoxytabersonine [M+H]^+^ 367.2021 (top) and TeT16H + Te16OMT3 product at *m/z* 367.2029 (bottom). Correlation score (GNPS-like score) of 0.93.

**Figure S19:**
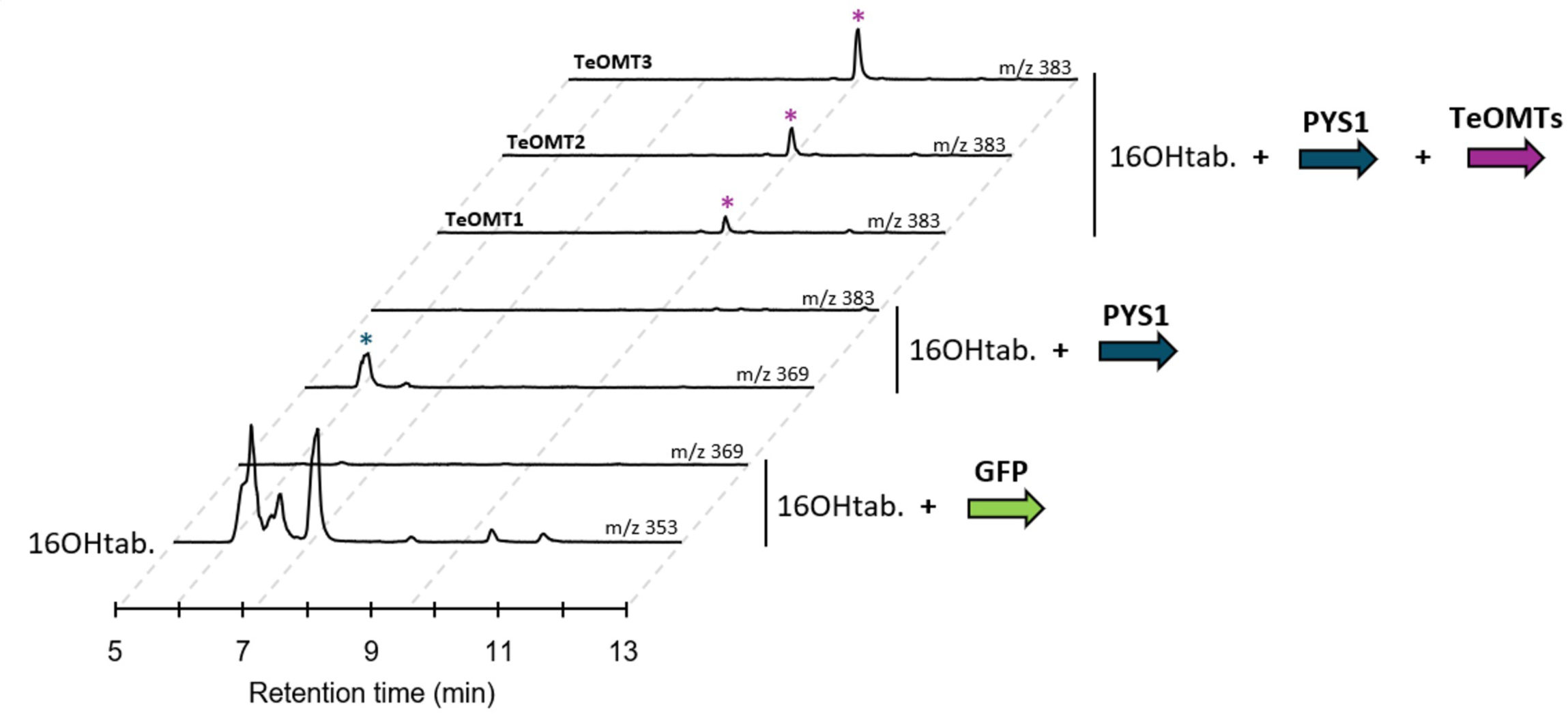
Biosynthesis of 16-methoxypachysiphine in tobacco by using the TeT16H, PYS1 and the three Te16OMTs. *PYS1*, the three characterized *Te16OMT* genes and *GFP* were transiently overexpressed in tobacco leaves in presence of 16-hydroxytabersonine produced by the TeT16H before extraction and concentration compound. Reaction products were analyzed by LC-MS. Positive differential mass chromatograms of each enzymatic step product are compared to GFP (negative control). Blue and purple asterisks indicate the 16-hydroxypachysiphine (*m/z* 369) and the 16-methoxypachysiphine (*m/z* 383), respectively. *16OHtab: 16-hydroxytabersonine*.

